# IDR-dependent MORC1 bodies limit chromatin accessibility to tune transposon repression

**DOI:** 10.64898/2025.12.14.694162

**Authors:** Yosuke Isota, Peilin Li, Kiichiro Sakata, Hiyu Chiwata, Hidetaka Kosako, Soichiro Yamanaka, Hiroya Yamazaki, Mikiko C. Siomi

## Abstract

Germline-specific MORC1-mediated transposon repression intersects with the MIWI2-piRNA pathway and involves H3K9me3 deposition, yet the underlying mechanism remains unclear. Here, we show that MORC1 assembles nuclear bodies in gonocytes. Using a doxycycline-inducible MORC1 expression system in NIH3T3 cells, we find that condensate formation is confined to the nucleus via an IDR containing a bipartite NLS, and that TRIM28, a corepressor that recruits the H3K9 methyltransferase SETDB1, is enriched in these bodies through protein−protein interactions. An IDR-deficient MORC1 mutant fails to form nuclear condensates but retains TRIM28 binding. ChIP-seq analysis reveals that this mutant exhibits higher occupancy at heterochromatin-embedded transposons than wild-type MORC1, with a pronounced preference for evolutionarily young transposon subfamilies that strongly overlap endogenous MORC1 targets in gonocytes. Wild-type MORC1 shows weaker enrichment at these elements and instead exhibits a greater tendency to occupy open chromatin. Condensates formed by exogenous MORC1 expression in NIH3T3 cells are substantially larger than those observed in gonocytes. These findings support a model in which MORC1-dependent transposon repression is governed by physical engagement with TRIM28 and by IDR-dependent, yet quantitatively restrained, nuclear body formation, which limits MORC1 chromatin accessibility, curtails off-target binding, and concentrates repression on *bona fide* transposon targets.

## Introduction

Microrchidia family CW-type zinc finger (MORC) proteins are evolutionarily conserved across diverse organisms, from prokaryotes to eukaryotes (Iyer et al., 2008). In humans, five paralogs, MORC1-MORC5 have been identified (Koch et al., 2017). The mouse genome likewise encodes five paralogs, MORC1, MORC2a, MORC2b, MORC3, and MORC4, each associated with distinct biological functions (Pastor et al., 2014; Shi et al., 2018; Tencer et al., 2020; Desai et al., 2021; Lee et al., 2021). Among them, MORC2a is a chromatin remodeler with GHKL ATPase activity that acts as an effector of the HUSH complex, contributing to heterochromatin compaction and gene silencing (Tchasovnikarova et al., 2017). MORC2b regulates the meiotic program (Shi et al., 2018), while MORC3 contributes to transposon silencing in mouse embryonic stem cells (Desai et al., 2021). By contrast, the function of MORC4 remains poorly understood. MORC1 uniquely plays a fertility-specific role; its deficiency causing male infertility, whereas no detectable effects have been observed on female reproductive function (Watson et al., 1998; Inoue et al., 1999; Pastor et al., 2014; Uneme et al., 2024).

MORC1 is highly expressed in mouse gonocytes, fetal male germ cells present from embryonic day 13.5 (E13.5) to postnatal day 3 (P3) (Uneme et al., 2024). In these cells, global DNA demethylation first occurs, followed by *de novo* DNA methylation, which is established around E16.5. Failure to reestablish DNA methylation leads to persistent transposon activation, which disrupts meiosis by inducing DNA double-strand breaks (Zamudio et al., 2015). Transposon silencing in gonocytes also involves histone modifications, such as H3K9 trimethylation (H3K9me3), in addition to DNA methylation (Uneme et al., 2024). In mice, these epigenetic mechanisms are mediated primarily by two protein families: Krüppel-associated box domain zinc finger proteins (KRAB-ZFPs) and PIWI proteins (Carmell et al., 2007; Fukuda et al., 2022).

KRAB-ZFPs bind specific transposon-derived genomic sequences and recruit the scaffold protein tripartite motif-containing 28 (TRIM28), which facilitates chromatin remodeling via interaction with factors such as the H3K9 methyltransferase SETDB1, the *de novo* DNA methyltransferase DNMT3, and the NuRD complex that mediates histone H3K9 deacetylation (Schultz et al., 2001; Schultz et al., 2002; Quenneville et al., 2012). These modifiers establish repressive chromatin states and DNA methylation at transposon loci. The KRAB-ZFP family has rapidly expanded in mammals, likely in response to evolving transposon threats (Jacobs et al., 2014; Imbeault et al., 2017).

PIWI proteins, loaded with PIWI-interacting RNAs (piRNAs), assemble piRNA-induced silencing complexes (piRISCs) that target transposon transcripts via RNA-RNA base pairing (Iwasaki et al., 2015). In mice, two PIWI proteins, MILI (PIWIL2) and MIWI2 (PIWIL4), are active in gonocytes, functioning in the cytoplasm and nucleus, respectively (Aravin et al., 2007; Kuramochi-Miyagawa et al., 2008). MIWI cleaves transposon transcripts in the cytoplasm, whereas MIWI2 suppresses transposons through transcriptional silencing involving DNA methylation and histone modification (Manakov et al., 2015). When a new transposon invades the germline genome, novel piRNAs are generated against the new element (Ernst et al., 2017). Thus, like KRAB-ZFPs, the PIWI-piRNA pathway also evolves in response to newly emerging transposons, reflecting an ongoing arms race.

We recently showed that transposons derepressed in MORC1-deficient gonocytes largely overlap with those derepressed in MIWI2 mutants, suggesting that MORC1 contributes to MIWI2-dependent transposon silencing (Uneme et al., 2024). In parallel, H3K9me3 levels decline at transposon loci in MORC1-deficient germ cells, accompanied by chromatin opening. Because TRIM28 recruits SETDB1 to transposons to establish H3K9me3 (Fukuda et al., 2022), these findings position MORC1 functionally within this axis. However, detailed mechanistic analysis has been lacking, in part because MORC1 expression is restricted to fetal male germ cells.

In this study, we first immunostained mouse gonocytes with anti-MORC1 antibodies and found that MORC1 forms discrete nuclear condensates. Because detailed analysis of these structures is technically challenging in gonocytes, we established a doxycycline (Dox)-inducible MORC1 expression system in NIH3T3 cells. Although NIH3T3 cells lack PIWI proteins, they retain the core gene-regulatory machineries, providing a tractable platform for mechanistic analyses.

## Results

### MORC1 assembles nuclear condensates via IDR-mediated phase separation

To assess endogenous MORC1 behavior, we immunostained fetal male germ cells with anti-MORC1 antibodies. MORC1 was detected in mouse Vasa homolog (MVH)-positive cells (Figure 1A), where it formed discrete nuclear foci with an estimated area of 0.013 nm^2^ (median; Supplementary Figure S1A). We hereafter refer to these condensates as MORC1 bodies. MVH is a germ-cell-specific RNA helicase that is widely used as a marker for identifying germ cells (Fujiwara et al., 1994).

**Figure 1.**
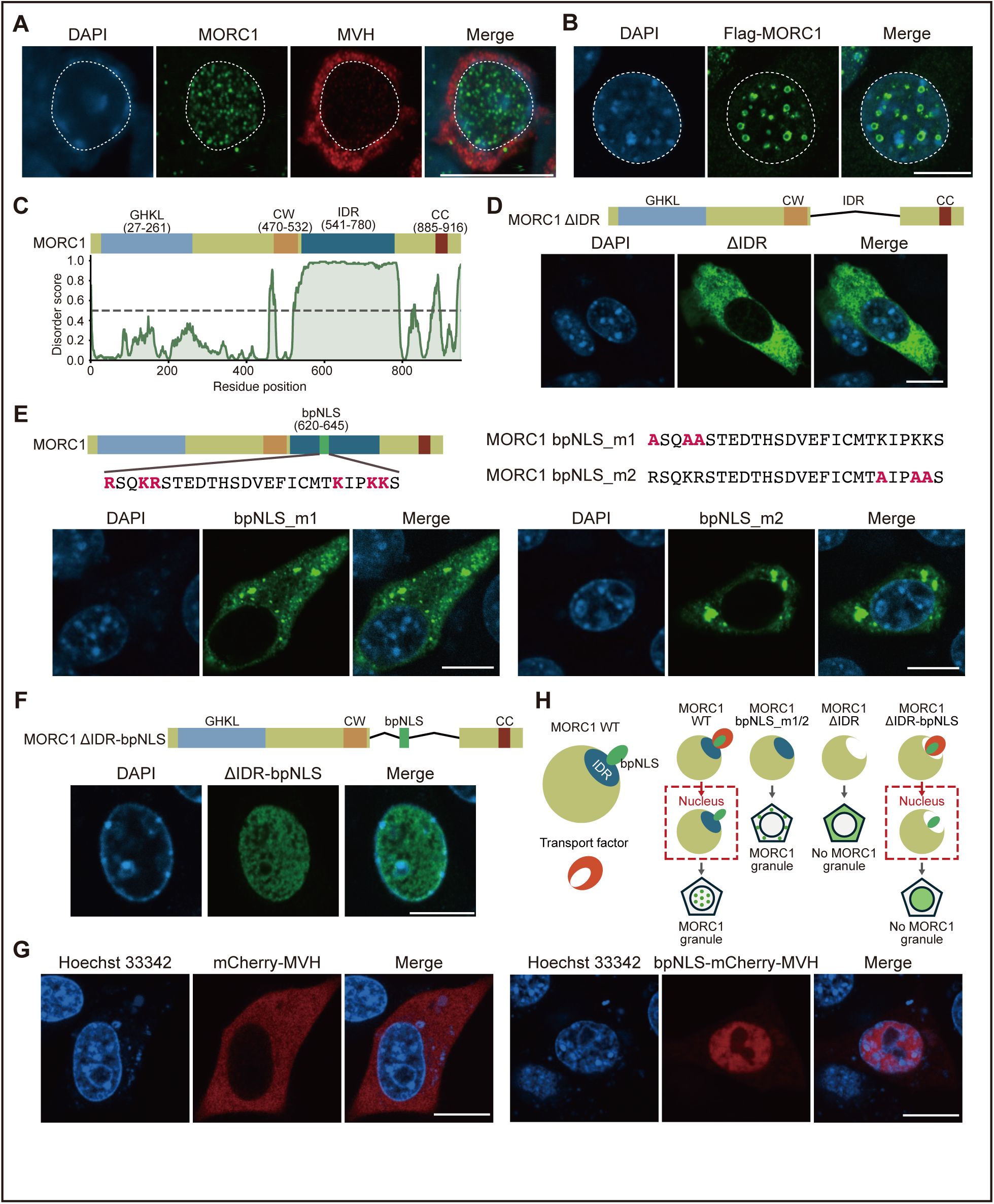
MORC1 assembles nuclear condensates through an IDR containing a bpNLS. **(A)** Endogenous MORC1 (green) forms discrete nuclear foci (“MORC1 bodies”) in MVH-positive (red) gonocytes. Nuclei (white dotted outline) are stained with DAPI (blue). Scale bar, 10 µm. **(B)** Upon Dox induction, Flag-MORC1 (green) forms nuclear condensates in NIH3T3 cells. Nuclei stained with DAPI (blue). Scale bar, 10 µm. **(C)** Schematic of MORC1 domain architecture and predicted IDR (blue, aa 541–780) immediately downstream of the CW domain (orange, aa 470–532). GHKL domain (gray, aa 27–261); coiled-coil (CC) domain (brown, aa 885–916). **(D)** MORC1 ΔIDR (green, ΔLeu541−Pro780), is dispersed throughout the cytoplasm in NIH3T3 cells. Nuclei stained with DAPI (blue). Scale bar, 10 µm. **(E)** bpNLS mutants (green) exhibit cytoplasmic localization. bpNLS_m1: Arg620, Lys623, and Arg624 were substituted with alanine; bpNLS_m2: Lys640, Lys643, and Lys644 were substituted with alanine. Nuclei stained with DAPI (blue). Scale bar, 10 µm. **(F)** The MORC1 ΔIDR-bpNLS mutant (green) localizes to the nucleus but fails to form condensates. Nuclei stained with DAPI (blue). Scale bar, 10 µm. **(G)** Fusion of the MORC1 bpNLS to mCherry-MVH (red) redirects the fusion protein to the nucleus. Nuclei stained with Hoechst 33342 (blue). Scale bar, 10 µm. **(H)** Model for MORC1 nuclear condensate assembly.

Because further multifaceted analysis of these condensates is technically challenging in gonocytes, we generated a Dox-inducible Flag-MORC1 line in NIH3T3 cells. Short-read RNA-seq (Wang et al., 2009) showed that NIH3T3 endogenously express *Morc2a* (NCBI gene ID: 74522), *Morc3* (338467), and *Morc4* (75746), but not *Morc1* (17450) or *Morc2b* (240069) (Supplementary Figures S1B and S1C). In contrast, RNA-seq data from gonocytes (Uneme et al., 2024) indicated high *Morc1* expression, followed by *Morc3* and *Morc2a*, with little or no *Morc2b* and *Morc4* (Supplementary Figure S1D).

Upon Dox induction, anti-Flag immunostaining revealed prominent MORC1 nuclear condensates that were substantially larger than MORC1 bodies in gonocytes (median area, 0.216 nm^2^) (Figure 1B and Supplementary Figure S1A). Signals were sometimes reduced in the condensate center, producing a ring-like morphology, consistent with antibody exclusion from dense phases or with proteins known to undergo phase separation (Pandey et al., 2025). Condensates frequently neighbored DAPI-bright structures (Figure 1B), likely constitutive heterochromatin or the nucleolus (Olins and Olins, 2005), although the functional relevance of this proximity remains unclear.

MORC1 bodies in gonocytes were much smaller and lacked a ring-like morphology (Supplementary Figure S1A), likely reflecting lower endogenous MORC1 levels and/or distinct nuclear architecture. Apparent attachment to DAPI-dense structures was not evident in gonocytes.

Because many condensate-forming proteins rely on intrinsically disordered regions (IDRs), we queried MORC1 with PrDOS (Ishida and Kinoshita, 2007). This identified a predicted IDR immediately C-terminal to the CW domain (Figure 1C and Supplementary Figure S1E). We deleted this segment and expressed the mutant (MORC1 ΔIDR) in NIH3T3 cells. In contrast to wild-type (WT) MORC1, the ΔIDR mutant failed to form nuclear foci and instead displayed a diffuse cytoplasmic distribution (Figure 1D), indicating that the IDR is required for nuclear condensate assembly and likely contributes nuclear localization.

### The MORC1 IDR harbors a bipartite NLS required for nuclear granule assembly and suppresses cytoplasmic granules

We queried the MORC1 IDR for nuclear localization signal(s) using NLS Mapper and identified a canonical bipartite nuclear localization signal (bpNLS) spanning Arg620–Ser645 (Figure 1E and Supplementary Figure S1E). The motif contains two basic clusters: Arg620–Arg624 near the N-terminal side and Lys640–Lys644 toward the C-terminal side.

To test function, we alanine-substituted three basic residues in either cluster, Arg620/Lys623/Arg624 (bpNLS_m1) or Lys640/Lys643/Lys644 (bpNLS_m2). Both mutants lost nuclear localization and instead formed cytoplasmic aggregates (Figure 1E), indicating that the predicted region serves as a *bona fide* bpNLS and that nuclear import, while necessary for nuclear granules, is not required for condensate assembly *per se*.

We next restored the bpNLS to the IDR-deleted mutant (ΔIDR-bpNLS). The fusion localized to nuclei but failed to assemble MORC1 granules (Figure 1F), showing that the IDR is indispensable for condensate formation, whereas the bpNLS is sufficient only for import. Fusing the MORC1 bpNLS to a predominantly cytoplasmic MVH (mCherry-MVH) drove its nuclear accumulation, whereas mCherry-MVH alone remained cytoplasmic (Figure 1G), demonstrating that the MORC1 bpNLS is sufficient to confer nuclear localization activity.

These results support a model (Figure 1H) whereby an import factor binds the MORC1 bpNLS to promote nuclear import and prevent cytoplasmic condensate formation; after import, its release permits IDR-driven granule assembly, confining condensates to the nucleus.

### MORC1 binds and concentrates TRIM28 into MORC1 granules

We profiled MORC1-interacting proteins in NIH3T3 cells. Following Dox-induced expression of Flag-MORC1, anti-Flag immunoprecipitates were analyzed by mass spectrometry (MS). Volcano plots from three biological replicates (Supplementary Figure S2A) revealed numerous proteins enriched in MORC1 immunoprecipitates (Figure 2A). To prioritize candidates, we overlaid STRING interaction scores using SETDB1, the H3K9me3 methyltransferase for transposon silencing in gonocytes as a network anchor (Liu et al., 2014; Cheng et al., 2021). TRIM28 emerged among the top hits with high STRING confidence (0.996) and strong MS enrichment (log2 abundance ratio > 3; −log10 P > 3.5) (Figures 2A and 2B). TRIM28 and additional network-supported factors are highlighted in Figures 2A and 2B. MORC1 robustly co-precipitated with TRIM28, whereas the negative control (mCherry) did not (Figure 2C), supporting the placement of MORC1 within the TRIM28/SETDB1 axis.

**Figure 2.**
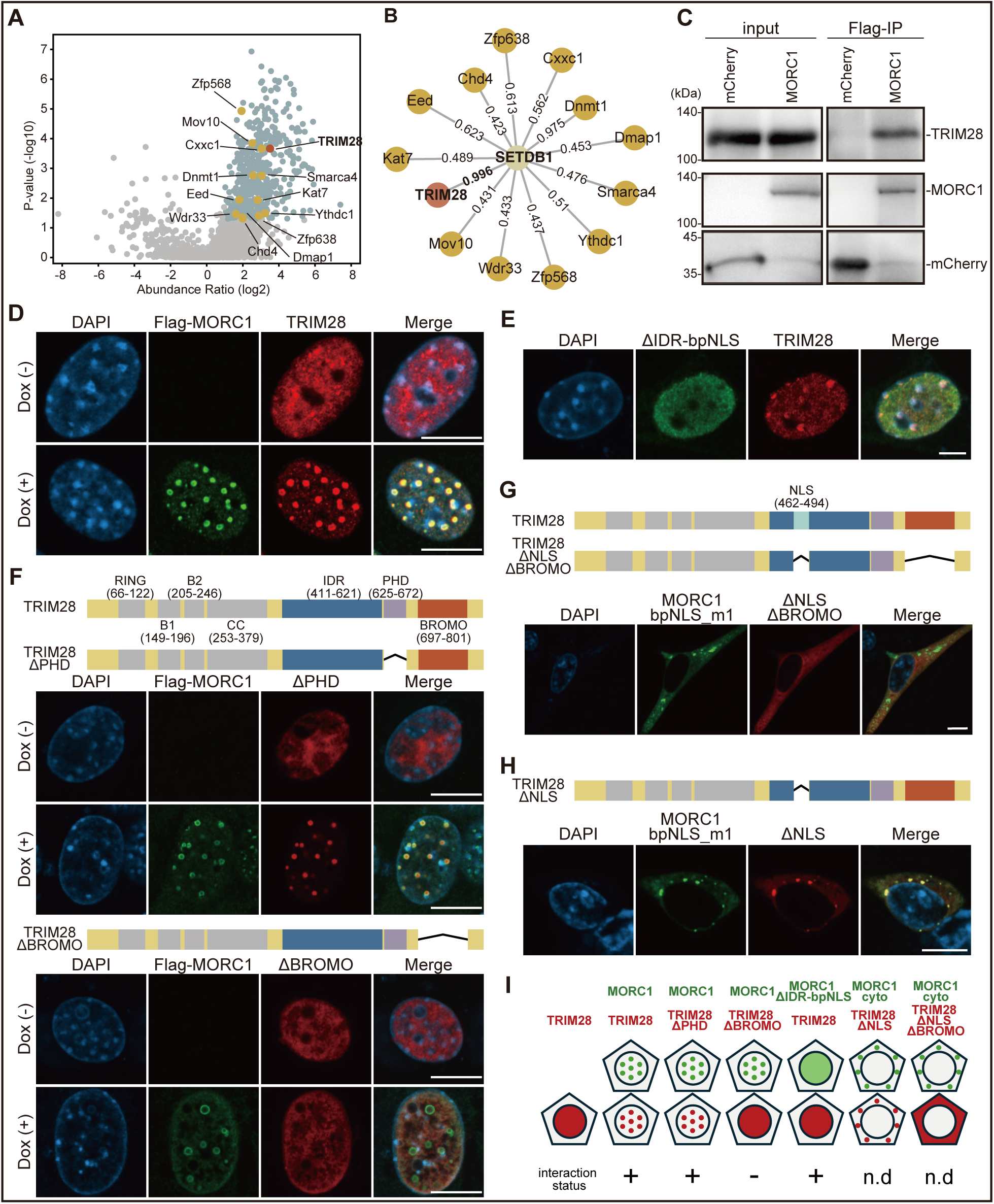
MORC1 binds and concentrates TRIM28 into MORC1 condensates. **(A)** Volcano plot showing proteins co-immunoprecipitating with MORC1 in NIH3T3 cells. Proteins appearing in (B) are specified by protein names. TRIM28 is shown in red. **(B)** STRING-based interaction network with SETDB1 as bait. **(C)** Co-immunoprecipitation of exogenously expressed MORC1 and TRIM28 in NIH3T3 cells. **(D)** TRIM28 (red) is diffusely distributed in the nucleoplasm of Dox (-) NIH3T3 cells but colocalizes with MORC1 (green) upon its expression. Nuclei stained with DAPI (blue). Scale bar, 10 µm. **(E)** TRIM28 (red) remains nucleoplasmic when MORC1 ΔIDR-bpNLS (green) is expressed instead of WT. Nuclei stained with DAPI (blue). Scale bar, 10 µm. **(F)** The TRIM28 ΔPHD mutant (red) is efficiently reposited into MORC1 condensates (green), whereas the ΔBROMO mutant (red) fails to localize to them. Nuclei stained with DAPI (blue). Scale bar, 10 µm. **(G)** TRIM28 ΔNLSΔBROMO (red) and MORC1-bpNLS_m1 (green) fail to colocalizes. Nuclei stained with DAPI (blue). Scale bar, 10 µm. **(H)** TRIM28 ΔNLS (red) and MORC1-bpNLS_m1 (green) colocalize in distinct cytoplasmic foci. Nuclei stained with DAPI (blue). Scale bar, 10 µm. **(I)** Summary of immunofluorescences.

Immunofluorescence showed that, in naïve NIH3T3 cells lacking MORC1 induction, endogenous TRIM28 was mainly nucleoplasmic [Dox (-); Figure 2D]. Upon MORC1 induction, TRIM28 sharply co-localized within MORC1 nuclear granules [Dox (+)], indicating that MORC1 recruits TRIM28 into MORC1 condensates through protein–protein interaction.

### TRIM28 bromodomain mediates TRIM28–MORC1 association

Replacing WT MORC1 with the IDR-deleted mutant (ΔIDR-bpNLS) abolished TRIM28 concentration: TRIM28 remained largely nucleoplasmic as in MORC1-deficient cells (Figure 2E). Notably, ΔIDR-bpNLS still co-immunoprecipitated endogenous TRIM28 (Supplementary Figure S2B), indicating that MORC1–TRIM28 binding is separable from MORC1 granule formation.

To delineate the TRIM28 region required for recruitment, we generated ΔPHD (PHD-domain deletion) and ΔBROMO (bromodomain deletion) mutants (Supplementary Figure S2C). The PHD domain promotes SUMOylation of the adjacent bromodomain, which then recruits repressors such as SETDB1 to silence targets, including transposons (Peng and Wysocka, 2008). Upon co-expression with WT MORC1, ΔPHD robustly concentrated into MORC1 granules, whereas ΔBROMO failed to localize to granules and remained diffusely nuclear (Figure 2F). Consistently, co-IP detected MORC1 interaction with ΔPHD but not with ΔBROMO (Supplementary Figure S2D), indicating that the bromodomain is essential for both MORC1 interaction and granule recruitment. TRIM28 also harbors a bpNLS within its IDR (Glu462–Asp494; Figure 2G and Supplementary Figure S2C). A ΔBROMO mutant lacking this NLS failed to co-aggregate with cytoplasmic MORC1-bpNLS_m1 (Figure 2G). However, restoring the bromodomain enabled a ΔNLS mutant to co-localize with MORC1-bpNLS_m1 (Figure 2H), further supporting the requirement for the bromodomain in MORC1-dependent granule recruitment. Figure 2I summarizes subcellular localization patterns from these immunofluorescence assays and the MORC1–TRIM28 interaction status.

### CW domain dictates family-specific TRIM28 concentration

Although MORC family members share similar architectures (Figure 3A), their behaviors in NIH3T3 cells diverged. MORC1, MORC3, and MORC4 formed nuclear foci (notably, MORC3 and MORC4 foci appeared larger), whereas MORC2a and MORC2b were predominantly nucleoplasmic (Figure 3B). Nuclear body formation has been reported for human MORC3 (Zhang et al., 2019). The nematode MORC-1 is implicated in nuclear RNAi and chromatin condensation (Weiser et al., 2017); however, to our knowledge, droplet-like condensates have not been directly demonstrated.

**Figure 3.**
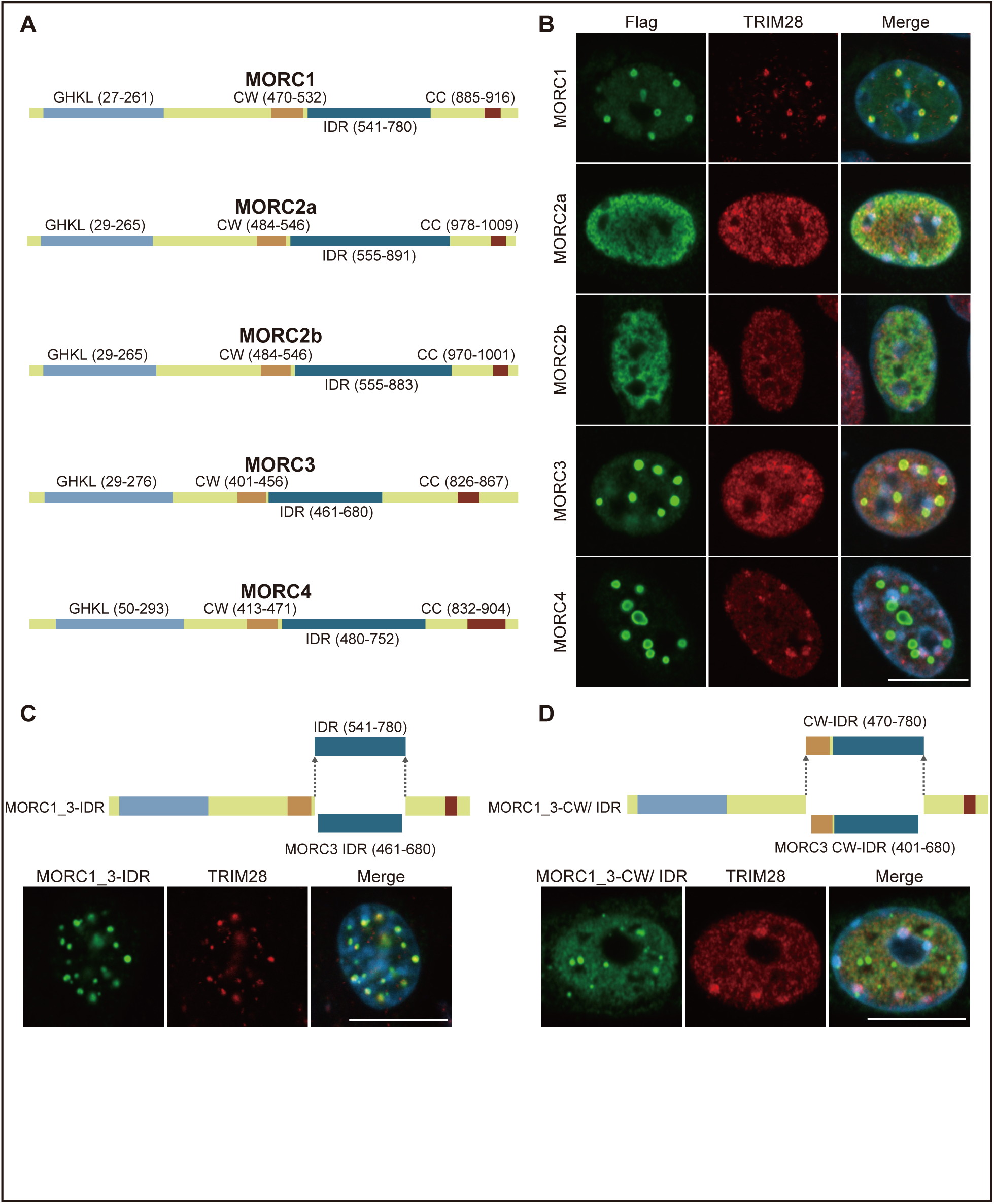
MORC1 exerts the strongest effect on the subcellular localization of TRIM28 among MORC family members. **(A)** Domain organization of MORC family members. The IDR of each member were determined based on their amino acid sequence homology with MORC1 IDR. **(B)** Subcellular localization of MORC1 family members (green) in NIH3T3 cells. TRIM28 is shown in red. Nuclei stained with DAPI (blue). Scale bar, 10 µm. **(C)** The MORC1_3-IDR chimera (green) concentrates TRIM28 (red) into nuclear condensates. Nuclei stained with DAPI (blue). Scale bar, 10 µm. **(D)** The MORC1_3-CW/IDR chimera (green) fails to concentrate TRIM28 (red) into the condensates. Nuclei stained with DAPI (blue). Scale bar, 10 µm.

We next compared how each MORC alters TRIM28 localization [Dox (-), Figure 2D]. MORC2a, MORC2b, and MORC4 had minimal effect; MORC3 partially shifted TRIM28 into foci; MORC1 produced the strongest accumulation (Figure 3B and Supplementary Figure S3A). Of the family, MORC2a, MORC3, and MORC4 are endogenously expressed in naïve NIH3T3 (Supplementary Figure S1C), consistent with their limited impact on TRIM28 (MORC3 modest).

Because the MORC1 IDR is required for granule formation, we asked whether domain differences explain functional divergence. Replacing the MORC1 IDR with that of MORC3 (MORC1_3-IDR) preserved TRIM28 reposition, comparable to WT MORC1 (Figure 3C). In contrast, swapping both the CW domain and IDR (MORC1_3-CW/IDR) markedly reduced concentration, phenocopying WT MORC3 despite robust nuclear foci (Figure 3D). The CW zinc-finger is a chromatin reader (He et al., 2010; Hoppmann et al., 2011); sequence identity between MORC1 and MORC3 CWs is low (31.7%; Supplementary Figure S3B), and the MORC3 CW preferentially binds H3K4me3, a mark of active genes (Guenther et al., 2007; Okitsu et al., 2010).

These data indicate that while IDR licenses condensate formation, the CW domain is the principal determinant of family specificity and governs interactions with other modules. Swapping in the CW domain of MORC3 retargets MORC1 to H3K4me3 marked chromatin while concomitantly weakening its interaction with TRIM28. This accounts for the attenuated TRIM28 recruitment observed with MORC1_3-CW/IDR. In this context, active marker guided chromatin recognition either competes with or dilutes TRIM28 binding.

### IDR-dependent control of MORC1 genomic occupancy

Nuclear co-localization of MORC1 and TRIM28, together with their protein−protein interaction, suggests that they may co-occupy genomic loci. To test this, we performed chromatin immunoprecipitation followed by sequencing (ChIP-seq) in NIH3T3 cells expressing Flag-MORC1 WT, using anti-Flag and anti-TRIM28 antibodies. Because the ΔIDR-bpNLS mutant failed to form nuclear condensates (Figure 1F) but retained TRIM28 binding at levels comparable to WT (Supplemental Figure 2B), we also analyzed this mutant (Figure 4A and Supplementary Figure S4A). To assess how the chromatin landscape changes in the presence or absence of MORC1, we additionally performed ATAC-seq in NIH3T3 cells expressing Flag-mCherry (control), Flag-MORC1 WT, or the ΔIDR-bpNLS mutant (Figure 4A and Supplementary Figure S4A).

**Figure 4.**
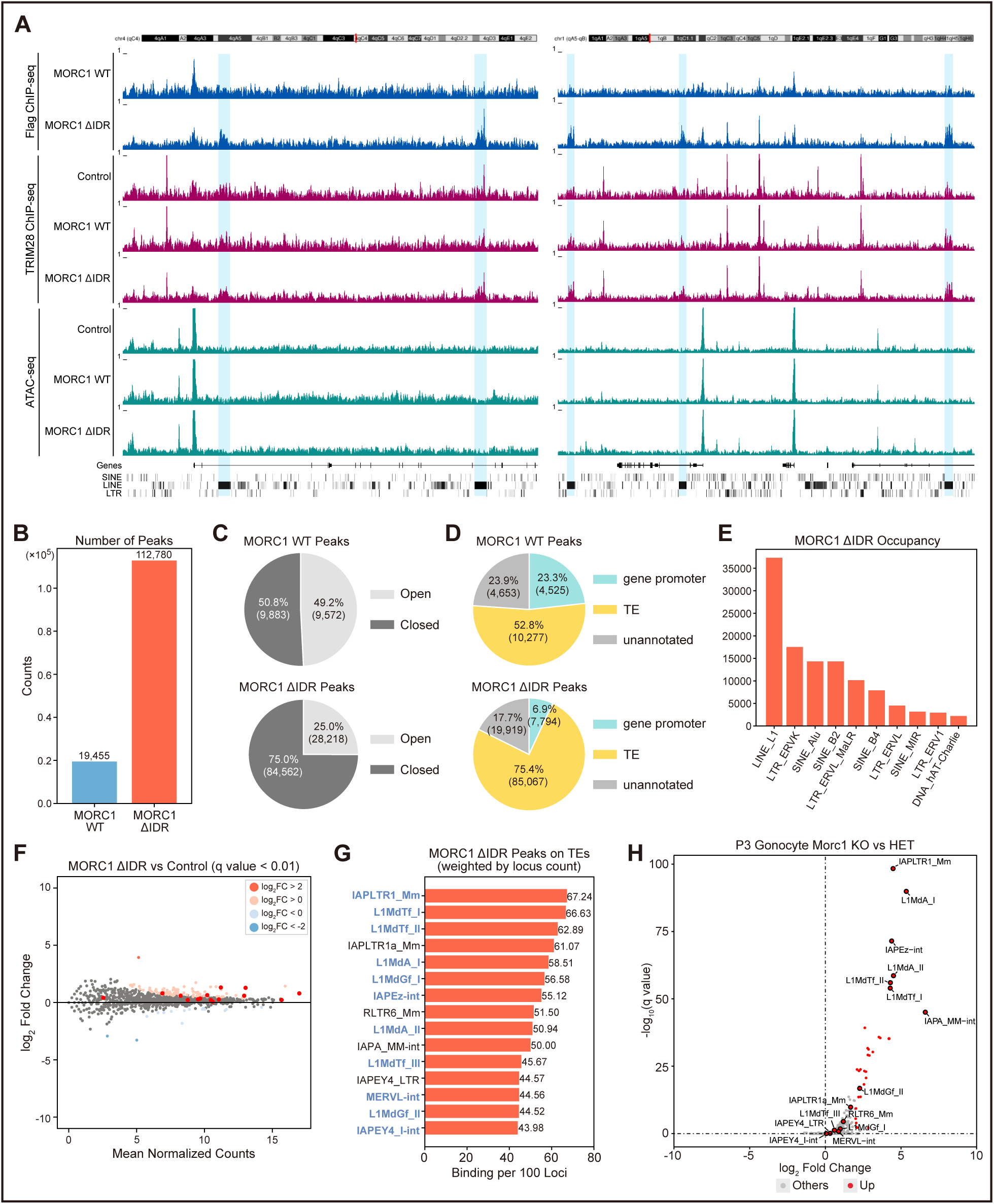
MORC1 preferentially associates with evolutionarily young transposons. **(A)** Genome browser view of MORC1 and TRIM28 ChIP–seq and ATAC–seq profiles. MORC1 ΔIDR indicates the MORC1 ΔIDR-bpNLS mutant, and mCherry was used as the control. Blue highlights mark de novo MORC1 peaks detected in ΔIDR compared with WT. **(B)** Number of MORC1 ChIP-seq peaks detected in WT (blue) and ΔIDR (red) samples. **(C)** Distribution of MORC1 ChIP-seq peaks in open or closed chromatin regions defined by ATAC-seq peaks. Upper, WT: lower, ΔIDR. **(D)** Genomic annotation of MORC1 ChIP-seq peaks as gene promoter, TE (transposon), or unannotated. Upper, WT: lower, ΔIDR. **(E)** Top 10 TE families most frequently occupied by MORC1 ΔIDR peaks. **(F)** MA-plot showing TE expression changes between NIH3T3 cells expressing MORC1 ΔIDR and mCherry (Control). **(G)** Top 15 transposon subfamilies most frequently occupied by MORC1 ΔIDR peaks; evolutionarily young subfamilies defined in Supplementary Figure 5(B) are highlighted in blue. **(H)** Volcano plot showing TE expression changes in P3 gonocytes from Morc1 KO and HET mice. Significantly upregulated TE subfamilies are highlighted in red, and subfamilies listed in (G) are annotated.

We first compared the ChIP-seq datasets for WT and mutant MORC1. Peak calling identified 19,445 peaks for WT and 112,780 peaks for the ΔIDR-bpNLS mutant (Figure 4B), even though WT and mutant MORC1 were expressed at comparable levels (Supplementary Figure S4B). Given that the principal difference between WT and mutant is the presence or absence of the IDR, we infer that IDR-dependent condensate formation, and the resulting increase in condensate volume, substantially influences MORC1’s genomic accessibility. This effect is particularly evident in NIH3T3 cells, where MORC1 condensates are markedly larger than those in fetal mouse germ cells (Figures 1A and 1B and Supplementary Figure S1A). These observations suggest that some factor(s) in gonocytes may constrain MORC1 body size, thereby limiting MORC1’s genomic accessibility and, consequently, the extent of its functional output. Differences in condensate size may also reflect differences between endogenous MORC1 levels in germ cells and exogenous MORC1 levels in NIH3T3 cells, although there is currently no way to directly compare, and therefore to adjust, these expression levels.

Comparison with ATAC-seq datasets revealed that 50.8% of WT peaks (*n* = 9,883) and 75.0% of mutant peaks (*n* = 84,562) fall within closed chromatin (Figure 4C). This result further suggests that IDR-driven condensate formation restricts MORC1 access to heterochromatic regions, likely because droplet formation efficiently increases in condensate volume of MORC1. However, contrary to our expectations, global chromatin accessibility (ATAC-seq) showed no major differences between WT or mutant MORC1 (Supplementary Figure S4C). Thus, in this context, exogenous MORC1 binding to NIH3T3 chromatin appears insufficient to remodel the pre-established chromatin landscape.

Almost all transposons in NIH3T3 cells reside in closed chromatin (98.3%; Supplementary Figure S4D). Together with the preferential localization of the ΔIDR-bpNLS mutant to closed chromatin (Figure 4C), this suggests that ΔIDR-bpNLS exhibits a stronger bias toward transposons than MORC1 WT. Consistent with this, 75.4% of ΔIDR-bpNLS peaks (*n* = 85,067) were located within transposon regions, whereas 52.8% of WT peaks (*n* = 10,277) fell within transposon regions (Figure 4D). We then counted the number of transposons that harbor WT or mutant MORC1 peaks. This analysis identified 17,508 and 119,105 transposons, respectively (Supplementary Figure S4E). Notably, 68.4% of transposons (*n* = 11,975) harboring WT MORC1 peaks were also occupied by mutant MORC1 peaks, indicating that MORC1’s overall preference for transposons is modestly influenced by the presence or absence of the IDR.

Although less prominent, MORC1, particularly WT, also showed appreciable binding to gene promoters (23.3%; Figure 4D). To identify which genes are preferentially targeted by MORC1 WT, we examined its promoter-bound genes and found enrichment for genes involved in RNA processing and/or translation, many of which are housekeeping genes (Supplementary Figure S4F). The mutant exhibited reduced promoter binding (6.9%; Figure 4D) but was associated with similar functional categories (Supplementary Figure S4G). This bias, regardless of IDR presence, suggests that MORC1 does not bind the genome in a purely stochastic manner. Promoter accessibility and expression of corresponding genes (by RNA-seq) were not appreciably altered by MORC1 binding (Supplementary Figure S5A). Thus, MORC1 binding to gene promoters alone appears to be insufficient to alter transcriptional output.

### MORC1 preferentially binds young LINE_1 and LTR_ERKV elements

The analyses so far suggested that, compared with WT, the ΔIDR-bpNLS mutant may more closely reflect the intrinsic genomic binding properties of MORC1 in gonocytes. Moreover, the number of MORC1-binding transposons was substantially higher in mutant cells (*n* = 119,105) than in WT cells (*n* = 17,508), corresponding to a 6.8-fold difference (Supplementary Figure S4E). In addition, WT binding largely overlapped with ΔIDR-bpNLS binding (68.4%; Supplementary Figure S4E). We therefore interpreted ΔIDR-bpNLS binding sites as “MORC1-mediated transposon binding events” and focused our subsequent analyses on these loci.

We first examined which classes of transposons MORC1 preferentially targets and observed a clear bias toward elements such as LINE_L1 and LTR_ERVK (Figure 4E), indicating that MORC1 is significantly enriched at specific transposon families. However, as with gene promoters, MORC1 binding to these transposons had little impact on their expression levels (Figure 4F). This is likely because, even in control NIH3T3 cells, these transposons are already embedded in heterochromatin (Supplementary Figure S4D) and are therefore repressed by endogenous silencing machineries.

We previously identified transposons that become derepressed upon loss of MORC1 in gonocytes (Uneme et al., 2024). Here, we compared these derepressed transposons with those bound by exogenously expressed MORC1 (ΔIDR-bpNLS) in NIH3T3 cells and found a striking overlap on evolutionarily young elements (Figures 4G and 4H, Supplementary Figure S5B). This suggests that MORC1, whether expressed endogenously in gonocytes or ectopically in NIH3T3 cells, tends to target a similar subset of transposons.

We also examined transposons bound by WT MORC1 (*n* = 17,508; Supplementary Figure S4E) but found only limited overlap with MORC1-repressed transposons in gonocytes (Supplementary Figure S5C). WT binding to these transposons had minimal impact on their expression levels (Supplementary Figure S5D). Together, these observations further support the notion that ΔIDR-bpNLS mutant more faithfully reflects the molecular function and genomic binding properties of MORC1 in gonocytes.

### MORC1 modulates TRIM28 recruitment to target transposons

We next analyzed the TRIM28 ChIP-seq datasets. We identified TRIM28 peaks embedded in transposons in control (mCherry) cells and in cells expressing MORC1 (ΔIDR-bpNLS) and then counted the number of associated transposons. In control cells, 67,977 transposons contained TRIM28 peaks, whereas in MORC1-expressing cells 140,788 transposons harbored TRIM28 peaks (Figure 5A), representing a 2.1-fold increase in TRIM28-targeted transposons upon expression of MORC1 (ΔIDR-bpNLS). These results indicate that even IDR-lacking MORC1, which fails to form nuclear condensates but retains its physical interaction with TRIM28, can enhance TRIM28 association with chromatin at transposon loci.

**Figure 5.**
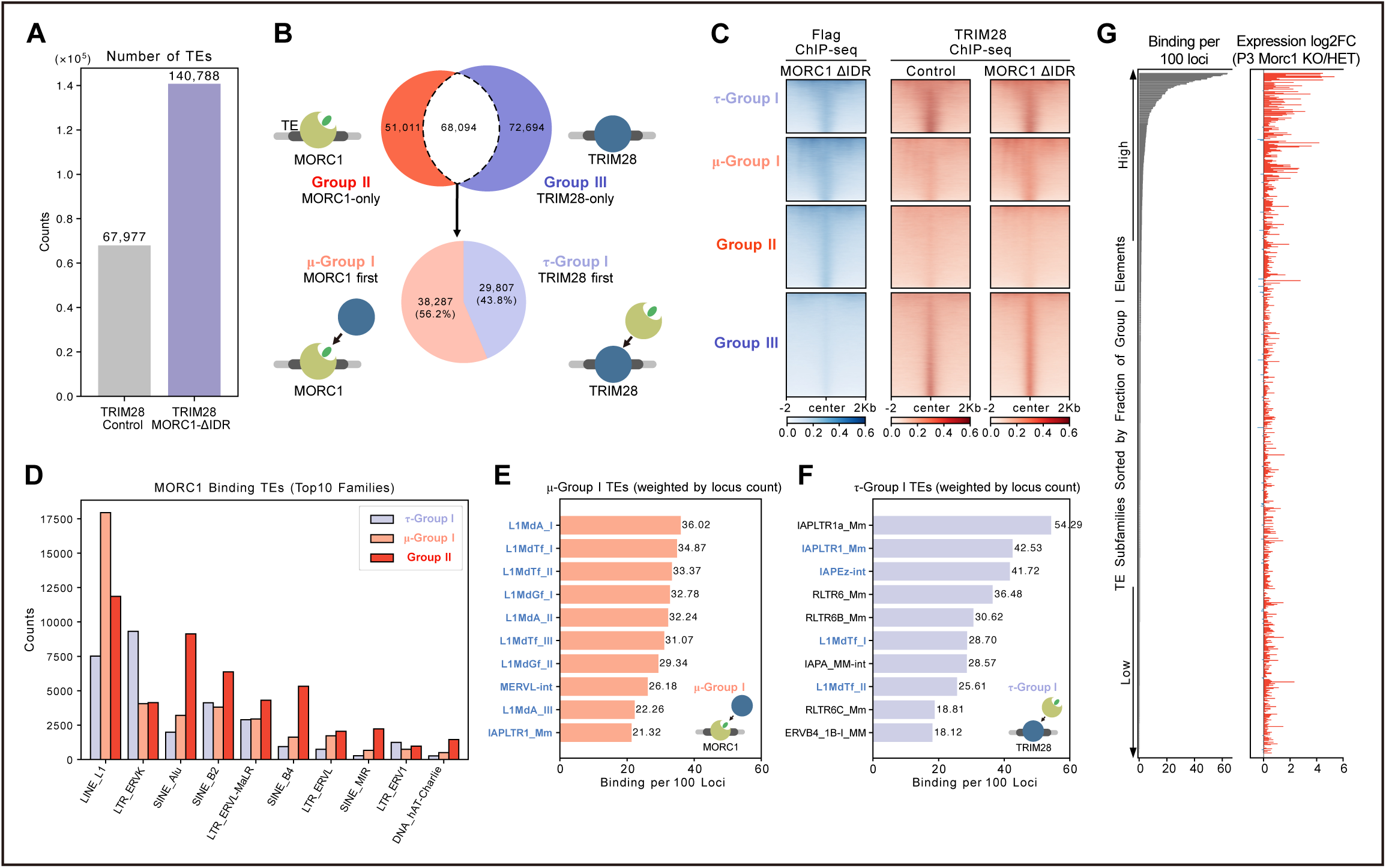
TRIM28 association with MORC ΔIDR at young transposon loci. **(A)** Number of TEs harboring TRIM28 ChIP-seq peaks in control (mCherry; gray) and MORC1 ΔIDR-expressing (purple) samples. **(B)** Classification scheme for TE harboring TRIM28 ChIP-seq peaks in the MORC1 ΔIDR sample (purple) in (A). Group I (white), transposons containing both TRIM28 and MORC1 ΔIDR peaks; Group II (red), MORC1 ΔIDR-only peaks, Group III (purple), TRIM28-only peaks. Group I transposons are further subdivided into μ-Group I (pale red), in which MORC1 engagement precedes TRIM28 (MORC1 first), and τ-Group I (pale purple), in which TRIM28 engagement precedes MORC1 (TRIM28 first). **(C)** Heatmaps showing MORC1 ΔIDR and TRIM28 enrichment across transposons in τ-Group I, μ-Group I, Group II, and Group III. MORC1 ΔIDR expression enhances TRIM28 accumulation at transposon loci of μ-Group I. **(D)** Transposon families most frequently represented among MORC1 binding elements (τ-Group I, μ-Group I, and Group II). **(E)** Top 10 transposon subfamilies within μ-Group I; evolutionarily young subfamilies defined in Supplementary Figure 5(B) are highlighted in blue. **(F)** Top 10 transposon subfamilies within τ-Group I; evolutionarily young subfamilies defined in Supplementary Figure 5(B) are highlighted in blue. **(G)** Correlation within TE subfamilies between the fraction of Group I (μ-Group I and τ-Group I) transposons (left) and expression changes in P3 MORC1-deficient gonocyes (right).

We next asked how many of these TRIM28-bound transposons were also associated with the MORC1 mutant. This analysis identified 68,094 transposons co-bound by TRIM28 and MORC1 (Group I, white area in Figure 5B). Among the remainder, 51,011 transposons were bound only by MORC1 (Group II, red), whereas 72,694 were bound only by TRIM28 (Group III, purple). We further subdivided Group I transposons into two categories: loci at which TRIM28 was already bound before MORC1 expression (TRIM28 first; τ-Group I) and loci at which TRIM28 binding appeared concomitantly with, or subsequently to, MORC1 binding (MORC1 first; μ-Group I). Of the Group I transposons, 43.8% (*n* = 29,807) fell into τ-Group I, whereas 56.2% (*n* = 38,287) were classified as μ-Group I (Figure 5B).

We also examined the distribution of MORC1 (ΔIDR-bpNLS) and TRIM28 across transposons in each group, i.e., τ-Group I, μ-Group I, Group II, and Group III (Figure 5C). MORC1 exhibited relatively stable enrichment on τ-Group I, μ-Group I, and Group II transposons, whereas, as expected, little enrichment was observed on Group III transposons. The heatmaps further revealed that TRIM28 was significantly recruited to transposons that were already occupied by MORC1 (μ-Group I; red panels in the second row from the top in Figure 5C). In contrast, only minor changes were observed in τ-Group I and Group II. In Group III, MORC1 expression led to increased TRIM28 accumulation at transposon loci, even though MORC1 itself was largely absent from these sites (Figure 5C). Together, these data clarify the distinct modes by which TRIM28 and MORC1 interact at transposon loci in NIH3T3 cells.

We next extracted MORC1-bound transposons from τ-Group I, μ-Group I, and Group II and classified them by transposon family. Each group displayed distinct features; for example, LINE_L1 elements were most abundant in μ-Group I, whereas LTR_ERVK elements predominated in τ-Group I (Figure 5D). To infer whether TRIM28 or MORC1 engages first at co-occupied loci, we quantified enrichment of individual transposon subfamilies within τ-Group I and μ-Group I. μ-Group I was markedly enriched for evolutionarily young LINE_L1 subfamilies (Figure 5E). In contrast, τ-Group I was biased toward LTR_ERVK elements, with particularly strong enrichment for IAP subfamilies (Figure 5F). Notable, transposons in both μ-Group I and τ-Group I showed a strong correspondence with those derepressed in MORC1-deficient gonocytes (Figure 5G), whereas this relationship was not evident for Group II (Supplementary Figure S5E and S5F). These results support a model in which MORC1, both in gonocytes and when ectopically expressed in NIH3T3 cells, preferentially associates with evolutionarily young transposons and promotes their repression in concert with TRIM28 and SETDB1.

## Discussion

In this study, using cultured NIH3T3 cells, we uncovered a previously unrecognized molecular mechanism underlying the functional deployment of MORC1, a gonocyte-specific transposon silencing factor. Because the cell number of fetal gonocytes is very low, its molecular characterization has been technically challenging. By establishing a Dox-inducible MORC1 expression system in NIH3T3 cells, we were able to bypass this limitation and begin to dissect its mode of action.

In NIH3T3 cells, MORC1 formed nuclear condensates resembling MORC1 bodies in gonocytes. Condensate formation was restricted to the nucleus via an IDR containing a bpNLS. Within the MORC family, MORC3 has been reported to form nuclear condensates in HeLa cells and to modulate transcriptional programs, in part by organizing PML nuclear bodies and recruiting factors such as p53 and CBP (Takahashi et al., 2007; Zhang et al., 2019). Whether and how other mouse MORC family members form similar condensates has only recently begun to be explored.

When we expressed each mouse MORC family member in NIH3T3 cells, MORC1, MORC3, and MORC4 formed condensates, whereas MORC2a and MORC2b did not and instead showed a diffuse nucleoplasmic distribution, despite all family members harboring predicted IDR(s). MORC2b is a rodent-specific retrotransposed derivative of MORC2a (Shi et al., 2018), consistent with their similar behavior. These paralogs may nonetheless form condensates in other cellular contexts, implying that condensate formation by MORC proteins may be highly context dependent.

TRIM28 is recruited to MORC1 granules through protein–protein interactions. Together with our previous bioinformatic analysis (Uneme et al., 2024), which indirectly linked the TRIM28/SETDB1 axis to MORC1, these data provide direct molecular support for a functional connection between these pathways. KRAB-ZFPs bind specific genomic sequences, recruit TRIM28 and SETDB1 to deposit H3K9me3, and repress transposon transcription (Wolf et al. 2015). In NIH3T3 cells, which normally lack MORC1; namely, KRAB-ZFP/TRIM28/SETDB1-mediated repression operates in a MORC1-independent manner, whereas in gonocytes, loss of MORC1 reduces H3K9me3 and leads to derepression of target transposons. Thus, the KRAB-ZFP/TRIM28/SETDB1 axis appears to engage heterochromatin through distinct, MORC1-dependent mechanisms in gonocytes compared with somatic cells.

The IDR-deleted MORC1 mutant (ΔIDR-bpNLS) failed to form nuclear granules yet retained TRIM28 binding. ChIP-seq analysis revealed that this mutant binds more strongly than WT MORC1 to transposons embedded in heterochromatin, particularly evolutionarily young transposon families that are targeted by MORC1 in gonocytes. In contrast, WT MORC1, whose molecular dynamics appears constrained by condensate formation, showed a tendency to occupy more open chromatin regions. Indeed, MORC1 condensates formed by exogenous MORC1 in NIH3T3 cells were markedly larger than MORC1 bodies in germ cells.

In summary, we propose a model in which MORC1 undergoes IDR-mediated condensate formation and, through protein–protein interactions, recruits TRIM28 into these condensates, thereby constraining the genomic search space and tuning the repressive activity of the TRIM28/SETDB1 axis at transposon loci in germ cells. Notably, WT MORC1 induces larger condensates in NIH3T3 cells than those observed in gonocytes and robustly accumulates TRIM28; however, it does not significantly reduce TRIM28 chromatin occupancy or otherwise inhibit TRIM28 function.

In gonocytes, MORC1 deficiency reduces H3K9me3 levels at transposon loci, suggesting that gonocyte-specific MORC1 functions as an intermediary scaffold that links TRIM28 and SETDB1 to their chromatin targets. Why such a higher-order regulatory layer is required, however, remains unclear. MORC1 is not expressed outside of gonocytes, raising the question of whether a functional counterpart operates in other cell types, or whether such a factor is dispensable there. These unsolved issues highlight the need for further investigation.

## Acknowledgments

We thank H. Ishizu for technical advice and experimental consultation and D. Nakazato, T. Fujisawa, and D. Brenske for experimental assistances. We also thank Y. Shinkai and all members of the Siomi laboratory for discussion. This work was supported by MEXT KAKENHI Grant Number JP25H01303 (to M.C.S.), JSPS KAKENHI Grant Numbers JP23K14141 and JP24H01271 (to H.Y.), Secom Science and Technology Foundation (to H.Y.), JSPS Program for Forming Japan’s Peak Research Universities (J-PEAKS) Grant Number JPJS00420240022 (to H.K.), Medical Research Center Initiative for High Depth Omics (to H.K.) and JST SPRING, Grant Number JPMJSP2108 (to P.L.).

## Author contributions

Y.I., H.Y., K.S., and H.C. performed biochemical experiments. H.K. performed LC-MS/MS analysis. P.L. performed bioinformatic analyses. S.Y. supervised and discussed the work. H.Y. and M.C.S. supervised and discussed the work and wrote the manuscript with the other authors.

## Materials and Methods

### Cell culture

NIH3T3 cells were cultured at 37°C with 5% CO_2_ in Dulbecco’s Modified Eagle Medium (Gibco) supplemented with 10% fetal bovine serum (Sigma) and Penicillin-Streptomycin-Glutamine (Gibco).

### Plasmid construction

Coding sequences for mouse MORC1, MORC2a, MORC2b, MORC3, MORC4, and MVH were amplified by PCR from either mouse testis cDNA or NIH3T3 cDNA library, depending on the gene. The TRIM28 open reading frame was obtained by gene synthesis (Invitrogen GeneArt). PCR products and linearized vector fragments were assembled into either pEGFPC1 (Abrisch et al., 2020) or Caspex (Myers et al., 2018) backbone by NEBuilder HiFi DNA Assembly Master Mix (New England Biolabs). This strategy was used to generate the MORC1 WT construct, MORC1 deletion derivatives, the MORC1 3-IDR and 3-CW/IDR variants, MORC paralog constructs, and the mCherry-or Flag-tagged MVH and TRIM28 WT expression plasmids. Site-directed mutagenesis for MORC1 bpNLS variants and TRIM28 domain-deletion constructs was performed by inverse PCR followed by Ligation high ver.2 (TOYOBO). Details of the PCR primers and plasmid constructs used in this study are provided in Supplementary Table 1.

### Plasmid transfection

NIH3T3 cells were seeded one day before to reach ∼70–90% confluency at the time of transfection. Plasmid DNA was diluted in Opti-MEM I Reduced Serum Medium (Gibco) and transfected using Lipofectamine 2000 (Thermo Fisher Scientific) according to the manufacturer’s instructions. For co-transfections, plasmids were mixed at equimolar ratios while keeping the total DNA amount constant. Medium was not exchanged after transfection. Cells were harvested or processed 48 h post-transfection for downstream analyses.

### Generation of doxycycline-inducible stable cell lines

Dox-inducible NIH3T3 cell lines expressing 3xFlag-mCherry, 3xFlag-MORC1 WT, or 3xFlag-MORC1 ΔIDR-bpNLS were generated using the piggyBac transposon system. Donor plasmids (Caspex-3xFlag-mCherry, Caspex-3xFlag-MORC1 WT, or Caspex-3xFlag-MORC1 ΔIDR-bpNLS) and the piggyBac transposase expression vector pCMV-hyPBase (Yusa et al., 2011) were co-transfected at a 5:1 (w/w) ratio. At 48 h post-transfection, cells were reseeded at approximately 1/21 of the original density. The following day, puromycin was added to the medium at 2 µg/mL for selection; medium was replaced every 3 days. Surviving colonies were picked individually and sequentially expanded from 24-well plates to 35 mm, 6 cm, and 10 cm dishes. Transgene expression was induced by addition of doxycycline (100 ng/mL) and cells were used for experiments 48 h after doxycycline addition.

### Co-immunoprecipitation

Cells were harvested by centrifugation and resuspended in hypotonic buffer [20 mM HEPES-KOH (pH 7.3), 10 mM KCl, 1.5 mM MgCl_2_, 0.5 mM DTT, 2 µg/mL pepstatin A, 2 µg/mL leupeptin, and 0.5 µg/mL aprotinin]. The suspension was pipetted and passed through a 25-gauge needle three times, then centrifuged; the pellet was washed three times with hypotonic buffer. The pellet was resuspended in co-IP buffer [50 mM HEPES-KOH (pH 7.3), 200 mM KCl, 1 mM EDTA, 1% Triton X-100, 0.1% sodium deoxycholate, 2 µg/mL pepstatin A, 2 µg/mL leupeptin, and 0.5 µg/mL aprotinin] and sonicated (Branson digital sonifier 250D-Advanced; amplitude 20%, 0.2 s ON / 0.8 s OFF cycles, total ON-time 1 min). Lysates were clarified by centrifugation at ∼20,000 × g for 20 min at 4°C. Dynabeads Protein G (Thermo Fisher Scientific) were washed with co-IP buffer and incubated with Flag M2 antibody (Sigma-Aldrich) for ≥30 min at room temperature with rotation to allow antibody binding. Clarified lysate was split and an input fraction reserved; the remainder was incubated with the antibody-bound beads for 2 h at 4°C with rotation. Beads were washed five times with co-IP buffer. Bound proteins were eluted in 2× SDS sample buffer [125 mM Tris-HCl (pH 6.8), 4% SDS, 19% glycerol, 200 mM DTT, and 0.01% bromophenol blue] at 70°C for 10 min prior to SDS-PAGE analysis.

### Western blotting and silver staining

Protein samples denatured in the sample buffer and boiled at 95℃ for 5 min. The proteins were separated by sodium dodecyl sulfate polyacrylamide electrophoresis using 8% polyacrylamide gels. After electrophoresis, they were transferred to ClearTrans Nitrocellulose Membrane (FUJIFILM Wako Pure Chemical). The membranes were blocked with 5% skim milk/PBS for 30 min at room temperature, primary antibodies diluted in Can Get Signal Solution 1 (TOYOBO) were applied overnight at 4℃. After washing, membranes were incubated with HRP-conjugated secondary antibodies diluted in Can Get Signal Solution 2 (TOYOBO) for 30 min at room temperature and signals detected using Clarity Western ECL Substrate (Bio-Rad) and imaged on ChemiDoc XRS Plus System (Bio-Rad). The following primary antibodies were used: anti-Flag (Sigma-Aldrich, 1:5,000 dilution), anti-β-Tubulin (Developmental Studies Hybridoma Bank, 1:1,000 dilution), anti-KAP1 (GeneTex, 1:1,000 dilution). The following secondary antibodies were used: anti-mouse IgG (H + L) HRP (Cappel, 1:5,000 dilution), anti-rabbit IgG HRP-linked (Cell Signaling Technology, 1:1,000 dilution). Silver staining was performed using SilverQuest Silver Staining Kit (Thermo Fisher Scientific) following the manufacturer’s instructions.

### Immunofluorescence of NIH3T3 cells

NIH3T3 cells were seeded on sterile glass coverslips placed in culture wells prior to transfection. At 48 h post-transfection, cells were fixed with methanol at room temperature for 5 min, followed by three washes with PBS containing 0.1% Tween-20 (PBS-T) for 10 min each. Cells were blocked with 1% BSA in PBS for 1 h at room temperature and incubated with primary antibodies diluted in 1% BSA/PBS for 1 h at room temperature. After three washes with PBS-T for 10 min each, cells were incubated with Alexa Fluor-conjugated secondary antibodies diluted in 1% BSA/PBS for 1 h at room temperature in the dark. Samples were washed three times with PBS-T and mounted with VECTASHIELD Antifade Mounting Medium with DAPI (VECTOR LABORATORIES). The following primary antibodies were used: anti-Flag (Sigma-Aldrich, 1:3,000 dilution), anti-KAP1 (GeneTex, 1:1,000 dilution). The following secondary antibodies were used: Alexa Fluor 488/555/594 goat anti-mouse IgG1 (Thermo Fisher Scientific, 1:1,000 dilution), Alexa Fluor 488 goat anti-rabbit IgG (H + L) (Thermo Fisher Scientific, 1:1,000 dilution).

### Immunofluorescence of testes

Mouse testes were fixed in 4% paraformaldehyde (pH 7.0) on ice for 1 h, washed four times with PBS, and cryoprotected sequentially in 10%, 20% and 30% sucrose in PBS at 4°C until tissues sank. Testes were embedded in O.C.T. Compound (Sakura Finetek Japan) and stored at −80°C. Cryosections (10 µm) were prepared and mounted onto glass slides. Sections were washed with Milli-Q water, subjected to antigen retrieval in 0.01 M citrate buffer (citric acid and trisodium citrate at a molar ratio of 1:4.6) at 105°C for 15 min, blocked with 3% skim milk in PBS-T for 30 min, and incubated with primary antibodies diluted in 3% skim milk/PBS-T for 1 h at room temperature. After three washes with PBS-T, sections were incubated with Alexa Fluor-conjugated secondary antibodies diluted in 3% skim milk/PBS-T for 1 h at room temperature in the dark. Slides were washed and mounted with VECTASHIELD Antifade Mounting Medium with DAPI (VECTOR LABORATORIES). The following primary antibodies were used: anti-MORC1 (Proteintech, 1:1,000 dilution), anti-MVH/DDX4 (Abcam 1:1,000 dilution). The following secondary antibodies were used: Alexa Fluor 488 goat anti-mouse IgG1 (Thermo Fisher Scientific, 1:1,000 dilution), Alexa Fluor 555 goat anti-rabbit IgG (H + L) (Thermo Fisher Scientific, 1:1,000 dilution).

### Live-cell imaging

NIH3T3 cells were cultured in glass-bottom dishes and transfected as described above. At 48 h post-transfection, nuclei were stained with Hoechst 33342 (Lonza) according to the manufacturer’s instructions. Live-cell imaging was performed using Stage Top Incubator (TOKAI HIT) to maintain cells at 37°C and 5% CO_2_ during image acquisition.

### RNA extraction and RNA-seq library construction

Total RNA was isolated from NIH3T3 cells using ISOGEN II (FUJIFILM Wako Pure Chemical) according to the manufacturer’s protocol. To remove residual genomic DNA, purified RNA was treated with Turbo DNase (Thermo Fisher Scientific) at 37°C for 30 min. DNase-treated RNA was then purified by phenol–chloroform extraction, followed by chloroform cleanup. The aqueous phase was subjected to ethanol precipitation with sodium acetate, washed with 70% ethanol, air-dried, and dissolved in nuclease-free water. RNA-seq libraries of wild-type NIH3T3 cells were prepared using TruSeq stranded mRNA (illumina) following manufacturer’s protocols and sequenced on an Illumina NovaSeq 6000 platform. RNA-seq libraries of NIH3T3 cells expressing 3xFlag-mCherry, 3xFlag-MORC1 WT, or 3xFlag-MORC1 ΔIDR-bpNLS were prepared using NEBNext Ultra II Directional RNA Library Prep Kit (New England Biolabs) following manufacturer’s protocols and sequenced on an Illumina NovaSeq X Plus platform.

### ChIP-seq library construction

Dox-inducible NIH3T3 cell lines expressing 3xFlag-mCherry, 3xFlag-MORC1 WT, or 3xFlag-MORC1 ΔIDR-bpNLS (treated with doxycycline for 24 h prior to the experiment, 1 × 10^6^) were fixed with 1% formaldehyde for 20 min. The cells were resuspended in Swelling buffer [20 mM Hepes (pH 7.9), 1.5 mM MgCl2, 10 mM KCl, 0.1% NP-40, and 1 mM DTT]. After incubation on ice for 20 min, pelleted nuclei were resuspended in 1× shearing buffer (Covaris) and fragmented to ∼500 bp with Branson digital sonifier SFX150 (BRANSON). Fragmented products diluted with RIPA buffer [50 mM Tris-HCl (pH 8.0), 150 mM NaCl, 2 mM EDTA (pH 8.0), 1% NP-40, 0.5% sodium deoxycholate, and 0.1% SDS] were mixed by rotation with Dynabeads Protein G (Thermo Fisher Scientific) for 1 h at 4℃. Immunoprecipitation was performed with 2 µL anti-Flag (Sigma-Aldrich), or 4 µL anti-KAP1 (GeneTex) on Dynabeads overnight at 4℃. Beads were washed with Low buffer [0.1% SDS, 1% Triton X-100, 2 mM EDTA (pH 8.0), 150 mM NaCl, and 20 mM Tris-HCl (pH 8.0)] three times and then with High buffer [0.1% SDS, 1% Triton X-100, 2 mM EDTA (pH 8.0), 500 mM NaCl, and 20 mM Tris-HCl (pH 8.0)] once. Immunoprecipitated products were eluted from beads by suspension in Direct elution buffer [10 mM Tris-HCl (pH 8.0), 5 mM EDTA (pH 8.0), 300 mM NaCl, and 0.5% SDS] while shaking at 65℃ for 15 min. The sample was treated with Proteinase K for 6 h at 37℃, followed by reversal of cross-linking overnight at 65℃. DNA was extracted by ethanol precipitation. Library preparation was carried out with a NEBNext Ultra Ⅱ DNA Library Prep Kit for Illumina (NEB) following the manufacturer’s instructions.

### ATAC-seq library construction

ATAC-seq was carried out following established protocols (Buenrostro et al., 2013; 2015), with minor adjustments for this study. Doxycycline-inducible NIH3T3 cells expressing 3×Flag-mCherry, 3×Flag-MORC1 (WT), or 3×Flag-MORC1 ΔIDR-bpNLS were collected after 48 h of doxycycline induction (1 × 10⁵ cells per reaction). Cells were gently lysed in 50 µL of ice-cold lysis buffer to release nuclei, which were subsequently pelleted and transferred into a transposition mixture composed of 2× TD buffer (25 µL; Illumina), TDE1 transposase (1.5 µL; Illumina), and nuclease-free water (23.5 µL). The transposition reaction proceeded at 37°C for 30 min. DNA fragments generated by TDE1 insertion were then purified using MinElute PCR Purification Kit (Qiagen). Libraries were produced by PCR amplification using barcoded primer sets (0.125 µM).

### Computational and image analysis

For the analysis of protein interactions, proteins demonstrating significant binding to MORC1 WT, in comparison to the control (444 proteins, log2Fold Change > 1 and P-value < 0.05), as well as SETDB1 (UniProt accession ID: O88974), were submitted to STRING (Version 12.0) (Szklarczyk et al., 2023). Subsequently, proteins interacting with SETDB1 were identified and extracted. The intrinsically disordered regions (IDRs) of MORC1 were predicted using DISOPRED3 (Jones and Cozzetto, 2015) via the PSIPRED server. Nuclear localization signals (NLSs) were predicted using cNLS Mapper (Kosugi et al., 2008; 2009a; 2009b) with a cut-off score of 5.0. For nuclear foci analysis, images of NIH3T3 cells and gonocytes expressing MORC1 WT were processed in Fiji (ImageJ, version 2.1.0/1.53c); nuclear foci were detected using the *Analyze Particles* function, and the area of each focus was measured.

### LC–MS/MS analysis

Proteins bound to the beads were digested with 200 ng of trypsin/Lys-C mix (Promega) at 37°C overnight. The resulting peptides were reduced, alkylated, acidified, and desalted using GL-Tip SDB (GL Sciences). Eluates were dried in a SpeedVac concentrator and reconstituted in 0.1% trifluoroacetic acid and 3% acetonitrile (ACN). LC-MS/MS analysis was performed on an EASY-nLC 1200 UHPLC system coupled to an Orbitrap Fusion mass spectrometer equipped with a nanoelectrospray ion source (Thermo Fisher Scientific). The peptides were separated on a 75 µm inner diameter × 150 mm C18 reversed-phase column (Nikkyo Technos) with a linear 4–32% ACN gradient for 0–100 min followed by an increase to 80% ACN for 10 min and a final hold at 80% ACN for 10 min. The mass spectrometer was operated in a data-dependent acquisition mode with a maximum duty cycle of 3 s. MS1 spectra were acquired with a resolution of 120,000, an automatic gain control (AGC) target of 4e5, and a mass range from 375 to 1,500 m/z. HCD MS/MS spectra were acquired in the linear ion trap with an AGC target of 1e4, an isolation window of 1.6 m/z, a maximum injection time of 35 ms, and a normalized collision energy of 30. Dynamic exclusion was set to 20 s. Raw data were searched against SwissProt database restricted to *Mus musculus* using Proteome Discoverer version 2.5 (Thermo Fisher Scientific, USA) with the Sequest HT search engine. The search parameters were as follows: (a) trypsin as an enzyme with up to two missed cleavages; (b) precursor mass tolerance of 10 ppm; (c) fragment mass tolerance of 0.6 Da; (d) carbamidomethylation of cysteine as a fixed modification; and (e) acetylation of protein N-terminus and oxidation of methionine as variable modifications. Peptides were filtered at a false discovery rate of 1% using the Percolator node. Label-free precursor ion quantification was performed using the Precursor Ions Quantifier node, and normalization was performed such that the total sum of abundance values for each sample over all peptides was the same.

### RNA-seq analysis

RNA-seq data from WT NIH3T3 cells were mapped to the mouse genome (mm10) using Bowtie2 in paired-end mode after quality check. Uniquely mapped reads were retained, while unmapped reads and reads overlapping genomic blacklist regions (mm10-blacklist.v2) were removed. Read counts on gene exons were then calculated using featureCounts with the options -p -t exon -g gene_id. RNA-seq data from Doxycycline-inducible NIH3T3 cells expressing 3×Flag-mCherry, 3×Flag-MORC1 WT, or 3×Flag-MORC1 ΔIDR-bpNLS were first adapter-trimmed using TrimGalore and processed using the pipeline described in a previous study (Teissandier et al., 2019). Specifically, trimmed reads were mapped to mouse genome (mm10) using STAR with following parameters - -runMode alignReads --outFilterMultimapNmax 5000 --outSAMmultNmax 1 --outFilterMismatchNmax 3 -outMultimapperOrder Random --winAnchorMultimapNmax 5000 --alignEndsType EndToEnd --alignIntronMax 1 --alignMatesGapMax 350 -- seedSearchStartLmax 30 --alignTranscriptsPerReadNmax 30000 -- alignWindowsPerReadNmax 30000 --alignTranscriptsPerWindowNmax 300 -- seedPerReadNmax 3000 --seedPerWindowNmax 300 --seedNoneLociPerWindow 1000. Read counts for transposons were summarized at the subfamily level using featureCounts with the options -p -M. Differentially expressed transposons between two conditions were identified using DESeq2 based on replicate count matrices. RNA-seq data of Morc1 KO and HET P3 gonocytes were analyzed with the same pipeline described above.

### ChIP-seq and ATAC-seq analysis

Adaptors of ChIP-seq reads were removed using Cutadapt, and reads were trimmed to 75 bp using SeqKit. Trimmed reads were aligned to the mouse genome (mm10) using Bowtie2 in paired-end mode. ATAC-seq reads were processed in the same way, with adaptor removal by Cutadapt followed by mapping to mm10 using Bowtie2 in paired-end mode with the options -N 1 -X 2000 --no-mixed --no-discordant. For aligned reads from both ChIP-seq and ATAC-seq, duplicates were removed using Picard, and only HighSignal regions within genomic blacklist regions (mm10-blacklist.v2) were removed. BAM files from biological replicates were merged, and bigWig files were generated using the deepTools function bamCoverage with CPM normalization. Peaks on the merged BAM files were identified using MACS3 with the parameters -f BAMPE --nolambda --keep-dup all. Enriched peaks for each sample were defined using the filterByExpr function in edgeR, with a minimum total count threshold of 10 for ChIP-seq peaks and 50 for ATAC-seq peaks. 21,846 protein-coding genes on autosomes and sex chromosomes were identified based on Mus_musculus.GRCm38.102.gtf from Ensemble (https://asia.ensembl.org/Mus_musculus/Info/Index) for quantitative analysis. Promoters were defined as ± 500 bp around the transcription start site (TSS). The mm10 transposon reference annotation was obtained from RepeatMasker (open-4.0.5), and entries annotated as Simple_repeat, Low_complexity, Satellite, rRNA, snRNA, tRNA, scRNA and other non-transposon categories were removed. Overlaps between gene promoters or transposons and peaks were identified using the intersect function in BEDTools. Genome-wide mappability was calculated with GenMap using the options -K 30 -E 2 -t -w, and the mean mappability for each transposon locus was obtained by intersecting the GenMap bedGraph with transposon coordinates using the map function in BEDTools.

## Supplementary Figure legends

**Supplementary Figure 1.**
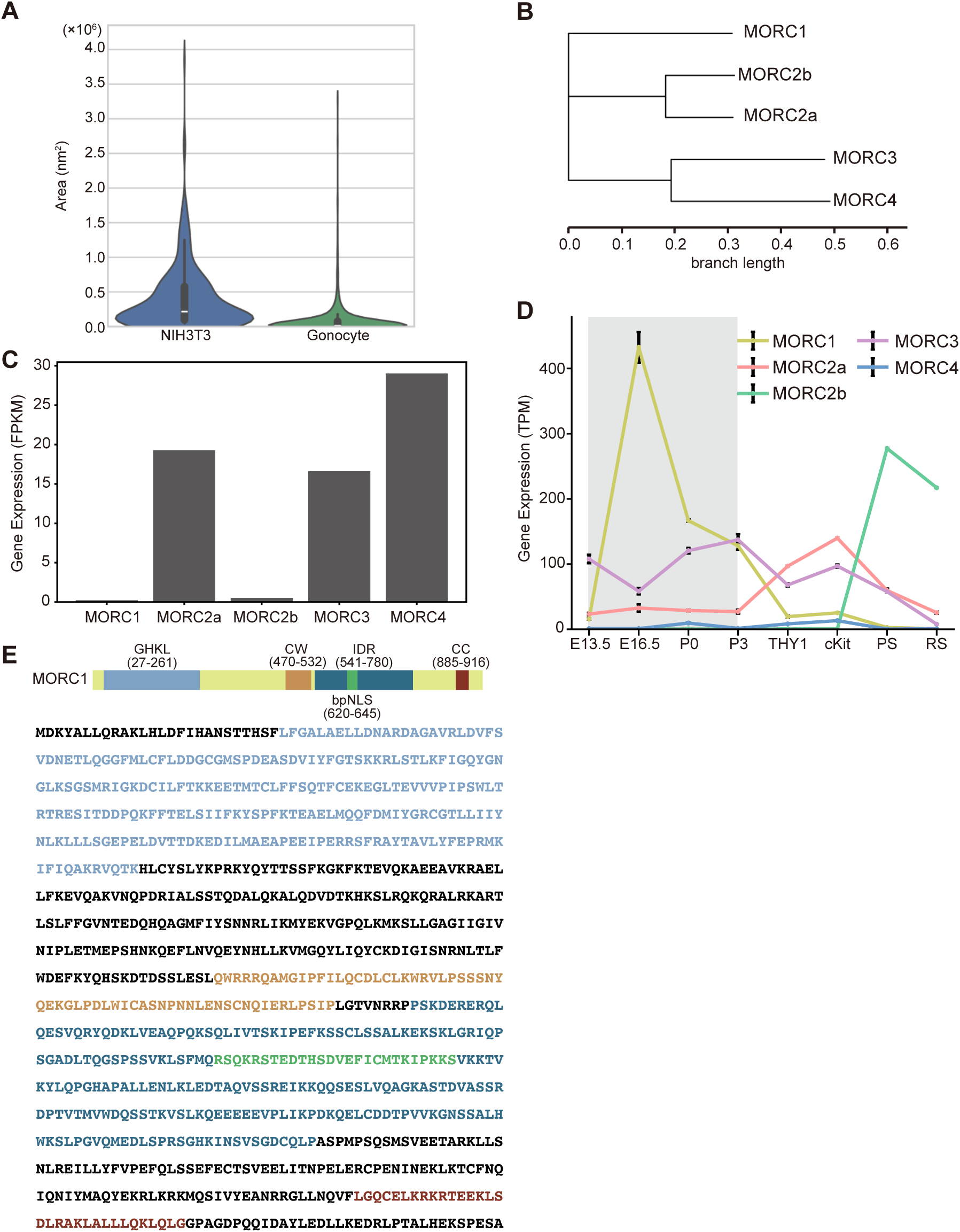
Mouse MORC family members. **(A)** Violin plots showing the distribution of MORC1 nuclear foci area in NIH3T3 cells and gonocytes. **(B)** Phylogenetic tree of mouse MORC family members (MORC1–4). Branch lengths represent the number of substitutions per site. **(C)** Bar plots showing the expression levels of MORC family members in NIH3T3 cells. **(D)** Line plots showing the expression levels of MORC family members during mouse male germ cell development, at embryonic stages E13.5 to P3 and in postnatal germ cell populations: THY1⁺ spermatogonia (undifferentiated SSCs), c-Kit⁺ spermatogonia (differentiating SSCs), pachytene spermatocytes (PS), and round spermatids (RS). **(E)** MORC1 peptide sequence and domains. GHKL (gray, aa 27−261); CW (orange, aa 470−532); IDR (blue, aa 541−780); bpNLS (green, aa 620−645); CC (brown, aa 885−916).

**Supplementary Figure 2.**
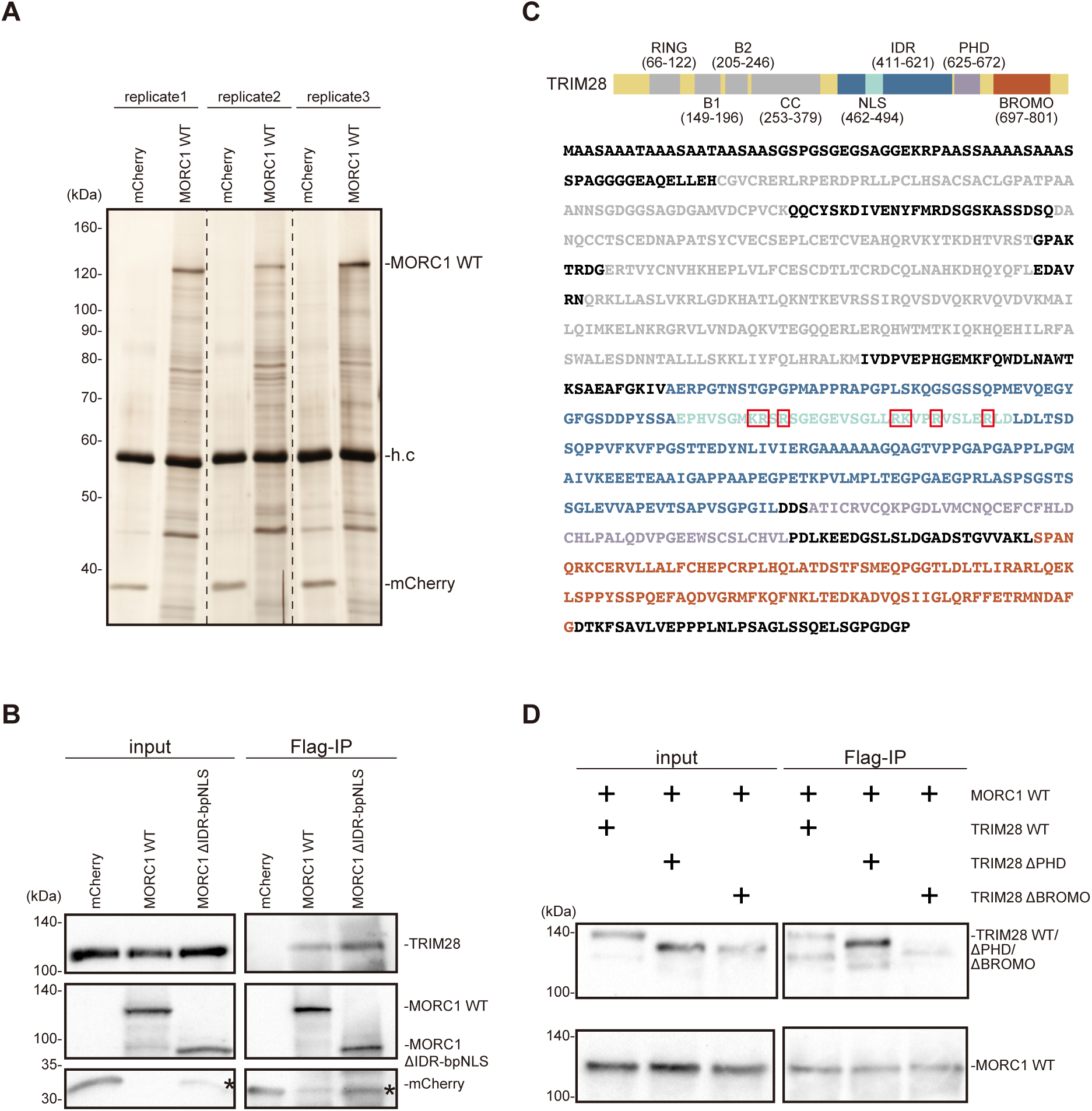
MORC1 interaction with TRIM28 and mapping of TRIM28 domains in NIH3T3 cells. **(A)** Silver-stained SDS-PAGE showing proteins immunoprecipitated with Flag antibody from NIH3T3 cells expressing 3xFlag-mCherry (control) or 3xFlag-MORC1 WT. **(B)** Co-immunoprecipitation of exogenously expressed MORC1 WT or ΔIDR-bpNLS and endogenously expressed TRIM28 in NIH3T3 cells. **(C)** Schematic representation of TRIM28 domains and the amino acid sequence, with sequence regions colored to match the domains in the schematic. Basic residues within the NLS (green) are highlighted with red boxes. **(D)** Co-immunoprecipitation of exogenously expressed MORC1 WT and exogenously expressed TRIM28 WT or ΔPHD or ΔBROMO in NIH3T3 cells.

**Supplementary Figure 3.**
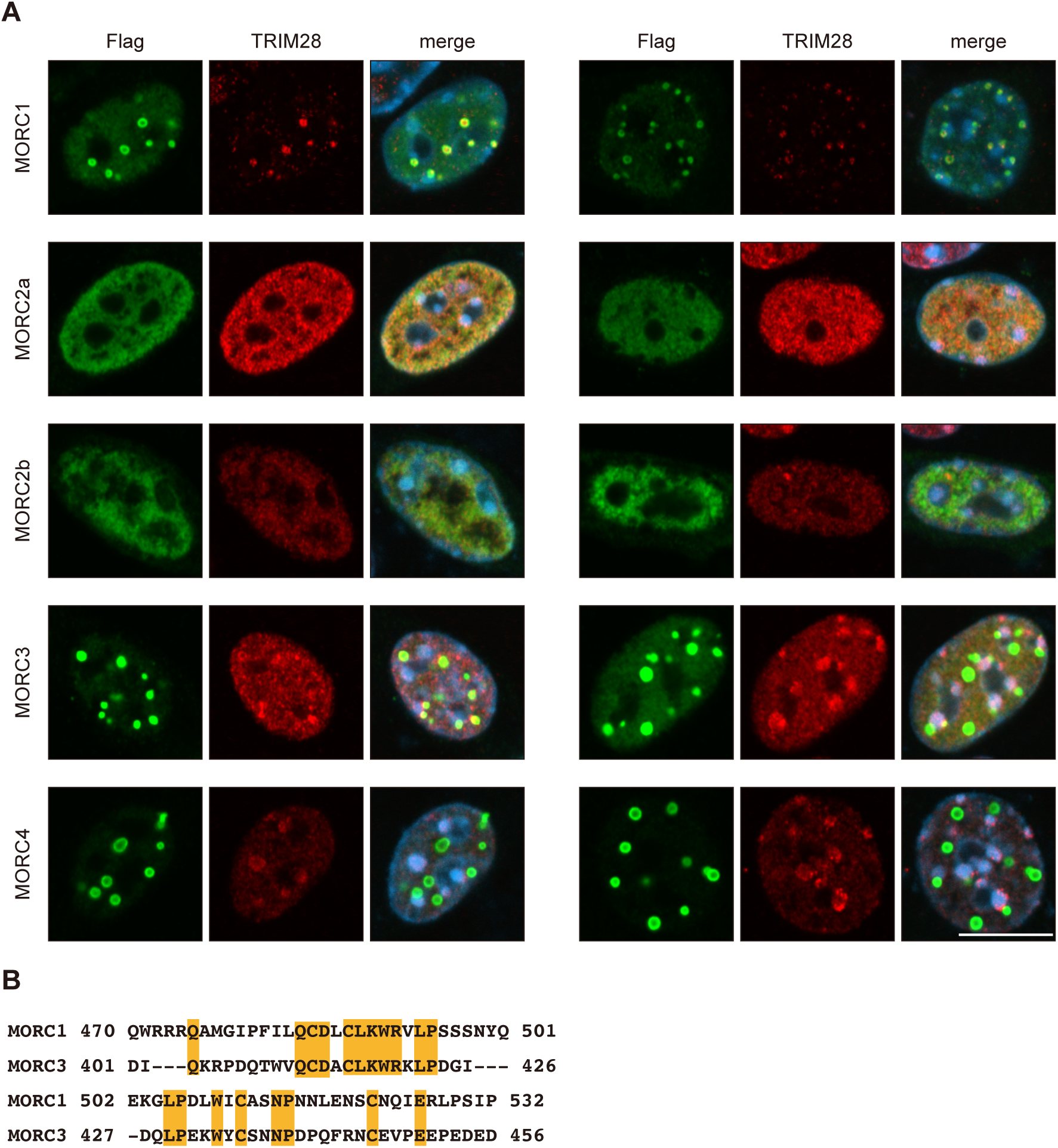
Subcellular localization of MORC family members and TRIM28 in NIH3T3 cells, and CW domain alignment between MORC1 and MORC3. **(A)** Subcellular localization of MORC family members in NIH3T3 cells (green). Subcellular localization of TRIM28 is also shown (red). Nuclei stained with DAPI (blue). Scale bar, 10 µm. **(B)** Amino acid sequence alignment of the CW domain from MORC1 and MORC3. Residues that are identical between the two proteins are highlighted in yellow.

**Supplementary Figure 4.**
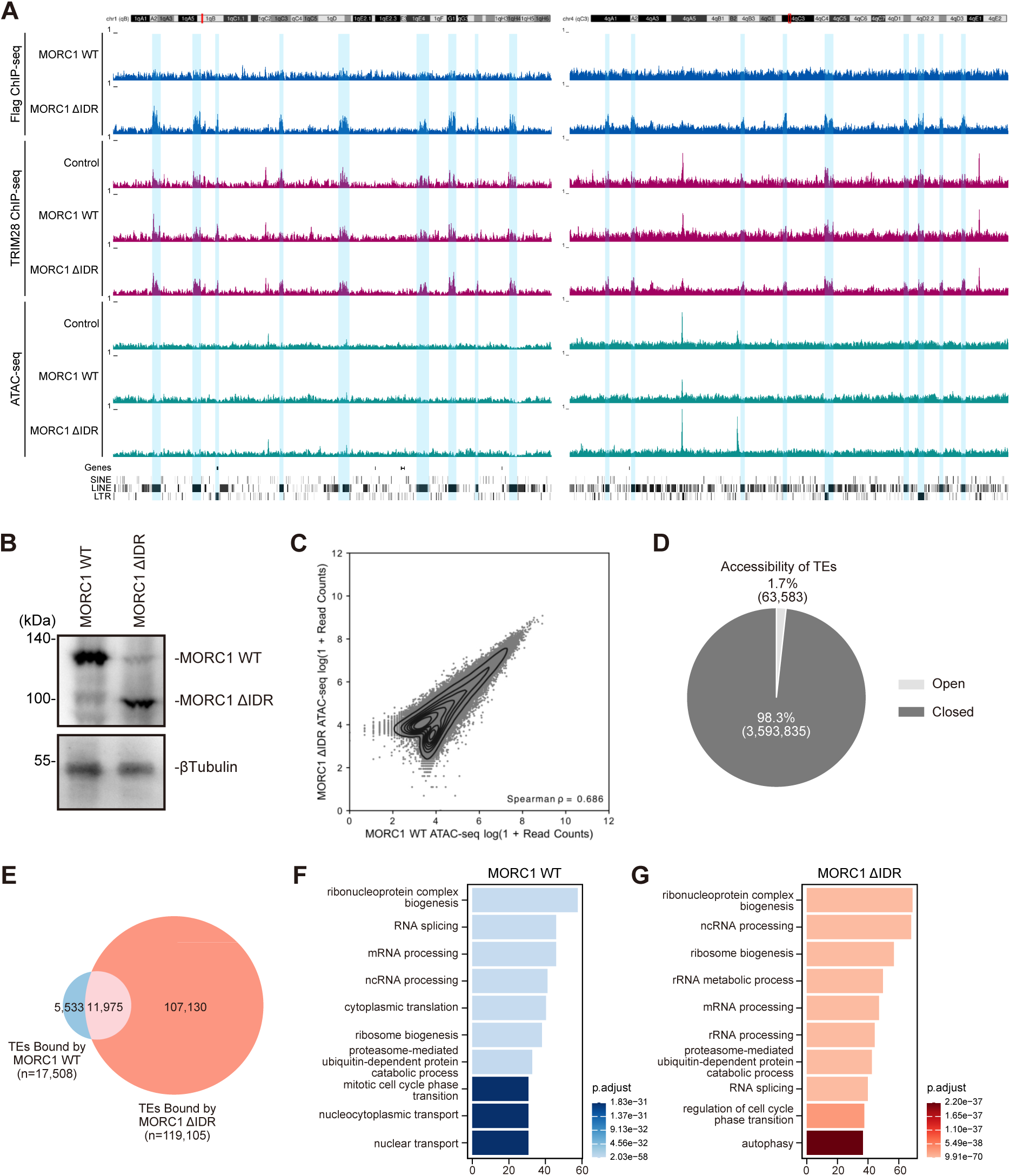
Genome-wide association and functional consequences of MORC1 WT and ΔIDR-bpNLS in NIH3T3 cells. **(A)** Genome browser view of MORC1 and TRIM28 ChIP–seq and ATAC–seq profiles. MORC1 ΔIDR indicates the MORC1 ΔIDR-bpNLS mutant, and mCherry was used as the control. Blue highlights mark de novo MORC1 peaks detected in ΔIDR compared with WT. **(B)** Western blot showing Dox-induced expression of MORC1 WT and ΔIDR in NIH3T3 cells. β-Tubulin, control. **(C)** Scatter plot comparing ATAC-seq read counts in universal peaks recognized in NIH3T3 cells expressing MORC1 WT (x-axis) and ΔIDR (y-axis). **(D)** Proportion of TEs classified as open or closed based on overlap with ATAC–seq peaks in NIH3T3 cells. **(E)** Venn diagram showing the number of TEs bound by MORC1 WT and ΔIDR. **(F) (G)** Bar plots showing Gene Ontology enrichment for genes with promoters bound by MORC1 WT (F) and MORC1 ΔIDR (G).

**Supplementary Figure 5.**
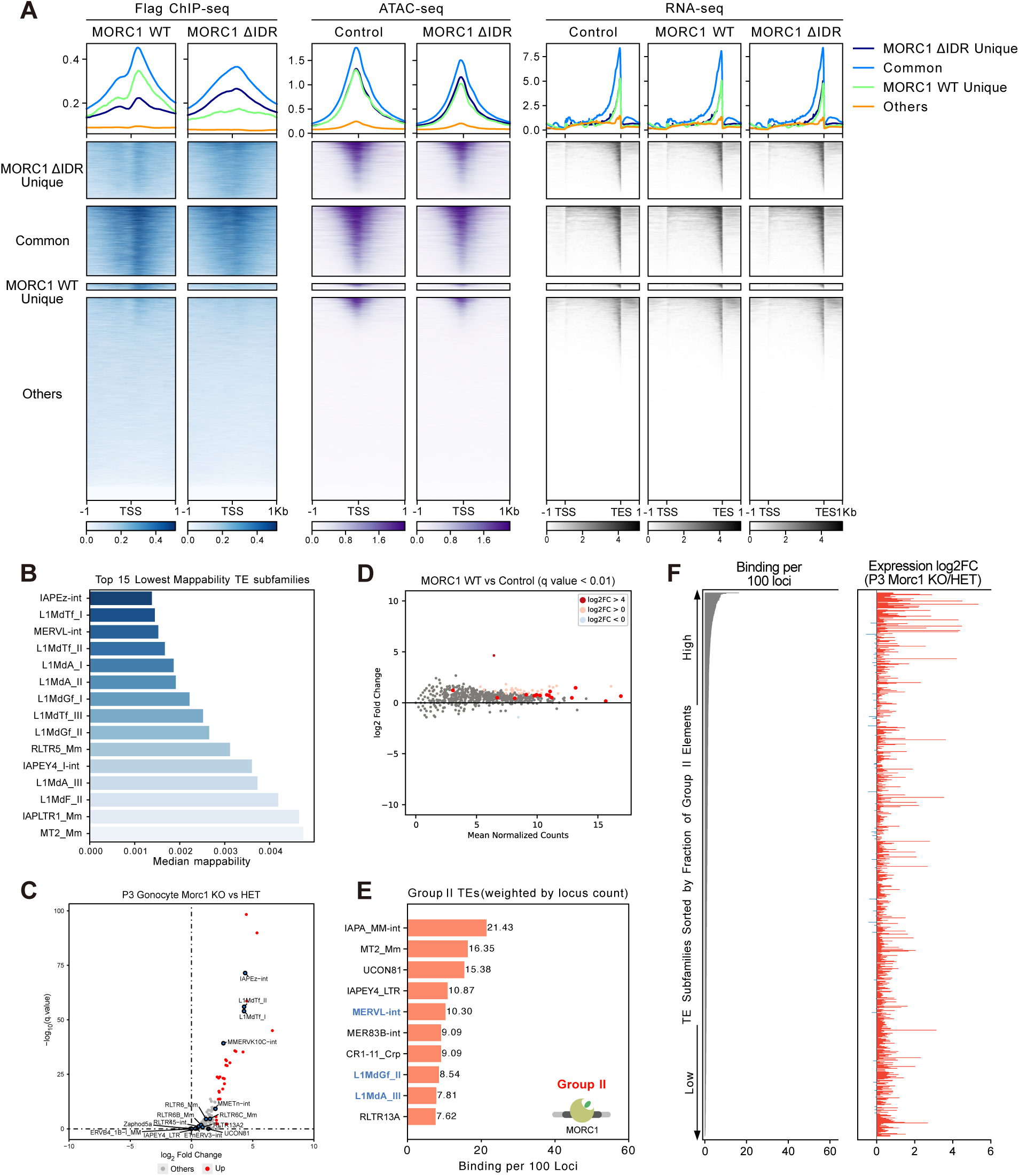
Genome-wide analyses of MORC1 binding and its impact on transposon regulation. **(A)** Heatmaps showing signals within promoters or entire gene regions in NIH3T3 cells. Left: flag ChIP-seq in MORC1 WT- and ΔIDR-expressing NIH3T3 cells. Middle: ATAC-seq in control (mCherry) and MORC1 ΔIDR-expressing NIH3T3 cells. Right: RNA-seq in control (mCherry), MORC1 WT-expressing, and MORC1 ΔIDR-expressing NIH3T3 cells. **(B)** Top 15 evolutionarily young TE subfamilies in mice, defined by the median mappability of each subfamily. **(C)** Volcano plot showing TE expression changes in P3 gonocytes from Morc1 KO and HET mice. Significantly upregulated TE subfamilies are highlighted in red, and subfamilies predominantly bound by MORC1 WT in NIH3T3 cells are annotated. **(D)** MA-plot showing TE expression changes between NIH3T3 cells expressing MORC1 WT and mCherry (Control). **(E)** Top 10 transposon subfamilies within Group II; evolutionarily young subfamilies as defined in (B) are highlighted in blue. **(F)** Correlation within TE subfamilies between the fraction of Group II elements (left) and expression changes in P3 MORC1-deficient gonocyes (right).

## References

Abrisch, Robert G., Samantha C. Gumbin, Brett Taylor Wisniewski, Laura L. Lackner, and Gia K. Voeltz. 2020. “Fission and Fusion Machineries Converge at ER Contact Sites to Regulate Mitochondrial Morphology.” Journal of Cell Biology 219 (4): e201911122. 10.1083/jcb.201911122.

Buenrostro, Jason D., Paul G. Giresi, Lisa C. Zaba, Howard Y. Chang, and William J. Greenleaf. 2013. “Transposition of Native Chromatin for Fast and Sensitive Epigenomic Profiling of Open Chromatin, DNA-Binding Proteins and Nucleosome Position.” Nature Methods 10 (12): 1213–18. 10.1038/nmeth.2688.

Buenrostro, Jason D., Beijing Wu, Howard Y. Chang, and William J. Greenleaf. 2015. “ATAC-Seq: A Method for Assaying Chromatin Accessibility Genome-Wide.” Current Protocols in Molecular Biology 109 (1): 21.29.1–21.29.9. 10.1002/0471142727.mb2129s109.

Carmell, Michelle A., Angélique Girard, Henk J. G. van de Kant, et al. 2007. “MIWI2 Is Essential for Spermatogenesis and Repression of Transposons in the Mouse Male Germline.” Developmental Cell 12 (4): 503–14. 10.1016/j.devcel.2007.03.001.

Cheng, Ee-Chun, Chia-Ling Hsieh, Na Liu, et al. 2021. “The Essential Function of SETDB1 in Homologous Chromosome Pairing and Synapsis during Meiosis.” Cell Reports 34 (1): 108575. 10.1016/j.celrep.2020.108575.

Desai, Varsha P., Jihed Chouaref, Haoyu Wu, et al. 2021. “The Role of MORC3 in Silencing Transposable Elements in Mouse Embryonic Stem Cells.” Epigenetics & Chromatin 14 (1): 49. 10.1186/s13072-021-00420-9.

Ernst, Christina, Duncan T. Odom, and Claudia Kutter. 2017. “The Emergence of piRNAs against Transposon Invasion to Preserve Mammalian Genome Integrity.” Nature Communications 8 (1): 1411. 10.1038/s41467-017-01049-7.

Fujiwara, Y, T Komiya, H Kawabata, et al. 1994. “Isolation of a DEAD-Family Protein Gene That Encodes a Murine Homolog of Drosophila Vasa and Its Specific Expression in Germ Cell Lineage.” Proceedings of the National Academy of Sciences 91 (25): 12258–62. 10.1073/pnas.91.25.12258.

Fukuda, Kei, Yoshinori Makino, Satoru Kaneko, et al. 2022. “Potential Role of KRAB-ZFP Binding and Transcriptional States on DNA Methylation of Retroelements in Human Male Germ Cells.” eLife 11 (March): e76822. 10.7554/eLife.76822.

Guenther, Matthew G., Stuart S. Levine, Laurie A. Boyer, Rudolf Jaenisch, and Richard A. Young. 2007. “A Chromatin Landmark and Transcription Initiation at Most Promoters in Human Cells.” Cell 130 (1): 77–88. 10.1016/j.cell.2007.05.042.

He, Fahu, Takashi Umehara, Kohei Saito, et al. 2010. “Structural Insight into the Zinc Finger CW Domain as a Histone Modification Reader.” Structure 18 (9): 1127–39. 10.1016/j.str.2010.06.012.

Hoppmann, Verena, Tage Thorstensen, Per Eugen Kristiansen, et al. 2011. “The CW Domain, a New Histone Recognition Module in Chromatin Proteins.” The EMBO Journal 30 (10): 1939–52. 10.1038/emboj.2011.108.

Imbeault, Michaël, Pierre-Yves Helleboid, and Didier Trono. 2017. “KRAB Zinc-Finger Proteins Contribute to the Evolution of Gene Regulatory Networks.” Nature 543 (7646): 550–54. 10.1038/nature21683.

Inoue, N., K. D. Hess, R. W. Moreadith, et al. 1999. “New Gene Family Defined by MORC, a Nuclear Protein Required for Mouse Spermatogenesis.” Human Molecular Genetics 8 (7): 1201–7. 10.1093/hmg/8.7.1201.

Ishida, Takashi, and Kengo Kinoshita. 2007. “PrDOS: Prediction of Disordered Protein Regions from Amino Acid Sequence.” Nucleic Acids Research 35 (Web Server issue): W460–464. 10.1093/nar/gkm363.

Iwasaki, Yuka W., Mikiko C. Siomi, and Haruhiko Siomi. 2015. “PIWI-Interacting RNA: Its Biogenesis and Functions.” Annual Review of Biochemistry 84: 405–33. 10.1146/annurev-biochem-060614-034258.

Iyer, Lakshminarayan M., Saraswathi Abhiman, and L. Aravind. 2008. “MutL Homologs in Restriction-Modification Systems and the Origin of Eukaryotic MORC ATPases.” Biology Direct 3 (March): 8. 10.1186/1745-6150-3-8.

Jones, David T., and Domenico Cozzetto. 2015. “DISOPRED3: Precise Disordered Region Predictions with Annotated Protein-Binding Activity.” Bioinformatics 31 (6): 857–63. 10.1093/bioinformatics/btu744.

Koch, Aline, Hong-Gu Kang, Jens Steinbrenner, D’Maris A. Dempsey, Daniel F. Klessig, and Karl-Heinz Kogel. 2017. “MORC Proteins: Novel Players in Plant and Animal Health.” Frontiers in Plant Science 8 (October). 10.3389/fpls.2017.01720.

Kosugi, Shunichi, Masako Hasebe, Tetsuyuki Entani, Seiji Takayama, Masaru Tomita, and Hiroshi Yanagawa. 2008. “Design of Peptide Inhibitors for the Importin α/β Nuclear Import Pathway by Activity-Based Profiling.” Chemistry & Biology 15 (9): 940–49. 10.1016/j.chembiol.2008.07.019.

Kosugi, Shunichi, Masako Hasebe, Nobutaka Matsumura, et al. 2009. “Six Classes of Nuclear Localization Signals Specific to Different Binding Grooves of Importin Α*.” Journal of Biological Chemistry 284 (1): 478–85. 10.1074/jbc.M807017200.

Kosugi, Shunichi, Masako Hasebe, Masaru Tomita, and Hiroshi Yanagawa. 2009. “Systematic Identification of Cell Cycle-Dependent Yeast Nucleocytoplasmic Shuttling Proteins by Prediction of Composite Motifs.” Proceedings of the National Academy of Sciences 106 (25): 10171–76. 10.1073/pnas.0900604106.

Kuramochi-Miyagawa, Satomi, Toshiaki Watanabe, Kengo Gotoh, et al. 2008. “DNA Methylation of Retrotransposon Genes Is Regulated by Piwi Family Members MILI and MIWI2 in Murine Fetal Testes.” Genes & Development 22 (7): 908–17. 10.1101/gad.1640708.

Lee, Geon Seong, Geon Kwak, Ji Hyun Bae, et al. 2021. “Morc2a p.S87L Mutant Mice Develop Peripheral and Central Neuropathies Associated with Neuronal DNA Damage and Apoptosis.” Disease Models & Mechanisms 14 (10): dmm049123. 10.1242/dmm.049123.

Liu, Sheng, Julie Brind’Amour, Mohammad M. Karimi, et al. 2014. “Setdb1 Is Required for Germline Development and Silencing of H3K9me3-Marked Endogenous Retroviruses in Primordial Germ Cells.” Genes & Development 28 (18): 2041–55. 10.1101/gad.244848.114.

Manakov, Sergei A., Dubravka Pezic, Georgi K. Marinov, William A. Pastor, Ravi Sachidanandam, and Alexei A. Aravin. 2015. “MIWI2 and MILI Have Differential Effects on piRNA Biogenesis and DNA Methylation.” Cell Reports 12 (8): 1234–43. 10.1016/j.celrep.2015.07.036.

Myers, Samuel A., Jason Wright, Ryan Peckner, Brian T. Kalish, Feng Zhang, and Steven A. Carr. 2018. “Discovery of Proteins Associated with a Predefined Genomic Locus via dCas9–APEX-Mediated Proximity Labeling.” Nature Methods 15 (6): 437–39. 10.1038/s41592-018-0007-1.

Okitsu, Cindy Yen, John Cheng Feng Hsieh, and Chih-Lin Hsieh. 2010. “Transcriptional Activity Affects the H3K4me3 Level and Distribution in the Coding Region.” Molecular and Cellular Biology 30 (12): 2933–46. 10.1128/MCB.01478-09.

Olins, Donald E, and Ada L Olins. 2005. “Granulocyte Heterochromatin: Defining the Epigenome.” BMC Cell Biology 6 (November): 39. 10.1186/1471-2121-6-39.

Pandey, Vikas, Tomohisa Hosokawa, Yasunori Hayashi, and Hidetoshi Urakubo. 2025. “Multiphasic Protein Condensation Governed by Shape and Valency.” Cell Reports 44 (4). 10.1016/j.celrep.2025.115504.

Pastor, William A., Hume Stroud, Kevin Nee, et al. 2014. “MORC1 Represses Transposable Elements in the Mouse Male Germline.” Nature Communications 5 (1): 5795. 10.1038/ncomms6795.

Peng, Jamy, and Joanna Wysocka. 2008. “It Takes a PHD to SUMO.” Trends in Biochemical Sciences 33 (5): 191–94. 10.1016/j.tibs.2008.02.003.

Quenneville, Simon, Priscilla Turelli, Karolina Bojkowska, et al. 2012. “The KRAB-ZFP/KAP1 System Contributes to the Early Embryonic Establishment of Site-Specific DNA Methylation Patterns Maintained during Development.” Cell Reports 2 (4): 766–73. 10.1016/j.celrep.2012.08.043.

Schultz, D. C., J. R. Friedman, and F. J. Rauscher. 2001. “Targeting Histone Deacetylase Complexes via KRAB-Zinc Finger Proteins: The PHD and Bromodomains of KAP-1 Form a Cooperative Unit That Recruits a Novel Isoform of the Mi-2alpha Subunit of NuRD.” Genes & Development 15 (4): 428–43. 10.1101/gad.869501.

Schultz, David C., Kasirajan Ayyanathan, Dmitri Negorev, Gerd G. Maul, and Frank J. Rauscher. 2002. “SETDB1: A Novel KAP-1-Associated Histone H3, Lysine 9-Specific Methyltransferase That Contributes to HP1-Mediated Silencing of Euchromatic Genes by KRAB Zinc-Finger Proteins.” Genes & Development 16 (8): 919–32. 10.1101/gad.973302.

Shi, Baolu, Jiangyang Xue, Jian Zhou, et al. 2018. “MORC2B Is Essential for Meiotic Progression and Fertility.” PLoS Genetics 14 (1): e1007175. 10.1371/journal.pgen.1007175.

Szklarczyk, Damian, Rebecca Kirsch, Mikaela Koutrouli, et al. 2023. “The STRING Database in 2023: Protein-Protein Association Networks and Functional Enrichment Analyses for Any Sequenced Genome of Interest.” Nucleic Acids Research 51 (D1): D638–46. 10.1093/nar/gkac1000.

Takahashi, Keiko, Naofumi Yoshida, Naoko Murakami, et al. 2007. “Dynamic Regulation of P53 Subnuclear Localization and Senescence by MORC3.” Molecular Biology of the Cell 18 (5): 1701–9. 10.1091/mbc.e06-08-0747.

Tchasovnikarova, Iva A., Richard T. Timms, Christopher H. Douse, et al. 2017. “Hyperactivation of HUSH Complex Function by Charcot–Marie–Tooth Disease Mutation in MORC2.” Nature Genetics 49 (7): 1035–44. 10.1038/ng.3878.

Teissandier, Aurélie, Nicolas Servant, Emmanuel Barillot, and Deborah Bourc’his. 2019. “Tools and Best Practices for Retrotransposon Analysis Using High-Throughput Sequencing Data.” Mobile DNA 10 (1): 52. 10.1186/s13100-019-0192-1.

Tencer, Adam H., Khan L. Cox, Gregory M. Wright, et al. 2020. “Molecular Mechanism of the MORC4 ATPase Activation.” Nature Communications 11 (1): 5466. 10.1038/s41467-020-19278-8.

Uneme, Yuta, Ryu Maeda, Gen Nakayama, et al. 2024. “Morc1 Reestablishes H3K9me3 Heterochromatin on piRNA-Targeted Transposons in Gonocytes.” Proceedings of the National Academy of Sciences 121 (13): e2317095121. 10.1073/pnas.2317095121.

Wang, Zhong, Mark Gerstein, and Michael Snyder. 2009. “RNA-Seq: A Revolutionary Tool for Transcriptomics.” Nature Reviews Genetics 10 (1): 57–63. 10.1038/nrg2484.

Watson, M. L., A. R. Zinn, N. Inoue, et al. 1998. “Identification of Morc (Microrchidia), a Mutation That Results in Arrest of Spermatogenesis at an Early Meiotic Stage in the Mouse.” Proceedings of the National Academy of Sciences of the United States of America 95 (24): 14361–66. 10.1073/pnas.95.24.14361.

Weiser, Natasha E., Danny X. Yang, Suhua Feng, et al. 2017. “MORC-1 Integrates Nuclear RNAi and Transgenerational Chromatin Architecture to Promote Germline Immortality.” Developmental Cell 41 (4): 408–423.e7. 10.1016/j.devcel.2017.04.023.

Wolf, Gernot, Peng Yang, Annette C. Füchtbauer, et al. 2015. “The KRAB Zinc Finger Protein ZFP809 Is Required to Initiate Epigenetic Silencing of Endogenous Retroviruses.” Genes & Development 29 (5): 538–54. 10.1101/gad.252767.114.

Zamudio, Natasha, Joan Barau, Aurélie Teissandier, et al. 2015. “DNA Methylation Restrains Transposons from Adopting a Chromatin Signature Permissive for Meiotic Recombination.” Genes & Development 29 (12): 1256–70. 10.1101/gad.257840.114.

Zhang, Yi, Bianca Bertulat, Adam H. Tencer, et al. 2019. “MORC3 Forms Nuclear Condensates through Phase Separation.” iScience 17 (July): 182–89. 10.1016/j.isci.2019.06.030.

Zhang, Yi, Brianna J. Klein, Khan L. Cox, et al. 2019. “Mechanism for Autoinhibition and Activation of the MORC3 ATPase.” Proceedings of the National Academy of Sciences of the United States of America 116 (13): 6111–19. 10.1073/pnas.1819524116.

